# The Structural Basis of the Genetic Code: Amino Acid Recognition by Aminoacyl-tRNA Synthetases

**DOI:** 10.1101/606459

**Authors:** Florian Kaiser, Sarah Krautwurst, Sebastian Salentin, V. Joachim Haupt, Christoph Leberecht, Sebastian Bittrich, Dirk Labudde, Michael Schroeder

## Abstract

Storage and directed transfer of information is the key requirement for the development of life. Yet any information stored on our genes is useless without its correct interpretation. The genetic code defines the rule set to decode this information. Aminoacyl-tRNA synthetases are at the heart of this process. For the first time, we extensively characterize how these enzymes distinguish all natural amino acids based on the computational analysis of crystallographic structure data. The results of this meta-analysis show that the correct read-out of genetic information is a delicate interplay between the composition of the binding site, non-covalent interactions, error correction mechanisms, and steric effects.

## Introduction

One of the most profound open questions in biology is how the genetic code was established. While proteins are encoded by nucleic acid blueprints, decoding this information in turn requires proteins. The emergence of this self-referencing system poses a chicken-or-egg dilemma and its origin is still heavily debated^1, 2^. Aminoacyl-tRNA synthetases (aaRSs) implement the correct assignment of amino acids to their codons and are thus inherently connected to the emergence of genetic coding. These enzymes link tRNA molecules with their amino acid cargo and are consequently vital for protein biosynthesis. Beside the correct recognition of tRNA features^3^, highly specific non-covalent interactions in the binding sites of aaRSs are required to correctly detect the designated amino acid^4–7^ and to prevent errors in biosynthesis^5, 8^. The minimization of such errors represents the utmost barrier for the development of biological complexity^9^ and accurate specification of aaRS binding sites is proposed to be one of the major determinants for the closure of the genetic code^10^. Beside binding side features, recognition fidelity is controlled by the ratio of concentrations of aaRSs and cognate tRNA molecules^11^ and may involve superordinate secondary structures in addition to side chain configurations^12, 13^.

### Evolution

The evolutionary origin of aaRSs is hard to track. Phylogenetic analyses of aaRS sequences show that they do not follow the standard model of life^14^; the development of aaRSs was nearly complete before the Last Universal Common Ancestor (LUCA)^15, 16^. Their complex evolutionary history included horizontal gene transfer, fusion, duplication, and recombination events^14, 17–21^. Sequence analyses^22^ and subsequent structure investigations^23, 24^ revealed that aaRSs can be divided into two distinct classes (*Class I* and *Class II*) that share no similarities at sequence or structure level. Each of the classes is responsible for 10 of the 20 proteinogenic amino acids and can be further grouped into subclasses^15^. Most eukaryotic genomes contain the complete set of 20 aaRSs. However, some species lack certain aaRS-encoding genes and compensate for this by post-modifications^7, 25–27^ or alternative pathways^28–30^. A scenario where Class I and Class II originated simultaneously from opposite strands of the same gene^31, 32^ is among the most popular explanations for the origin of aaRSs. This so-called Rodin-Ohno hypothesis (named after Sergei N. Rodin and Susumu Ohno^31^) is supported by experimental deconstructions of both aaRS classes^33–35^. At the dawn of life the concurrent duality could have allowed to implement an initial binary choice, which is the minimal requirement to establish any code^9^.

### Biochemical Function

In order to fulfill their biological function aaRSs are required to catalyze two distinct reaction steps. Prior to its covalent attachment to the 3’ end of the tRNA molecule, the designated amino acid is activated with adenosine triphosphate (ATP) and an aminoacyl-adenylate intermediate is formed^36, 37^. In general, the binding sites of aaRSs can be divided into two moieties: the part where ATP is bound as well as the part where specific interactions with the amino acid ligand are established (Fig. 1). Is is assumed that the amino acid activation with ATP constituted the principal kinetic barrier for the creation of peptides in the prebiotic context^35^. Due to the fundamental importance of this first reaction step, highly conserved sequence^4^ and structural motifs^38^ exist, which are likely to be vital for the aminoacylation reaction. While the activation of amino acids with ATP is the basic requirement of all aaRSs and is consistent within each aaRS class^38^, the recognition mechanism of individual amino acids differs substantially between each aaRS. These differences are among the key drivers to maintain a low error rate during the translational process.

**Figure 1.**
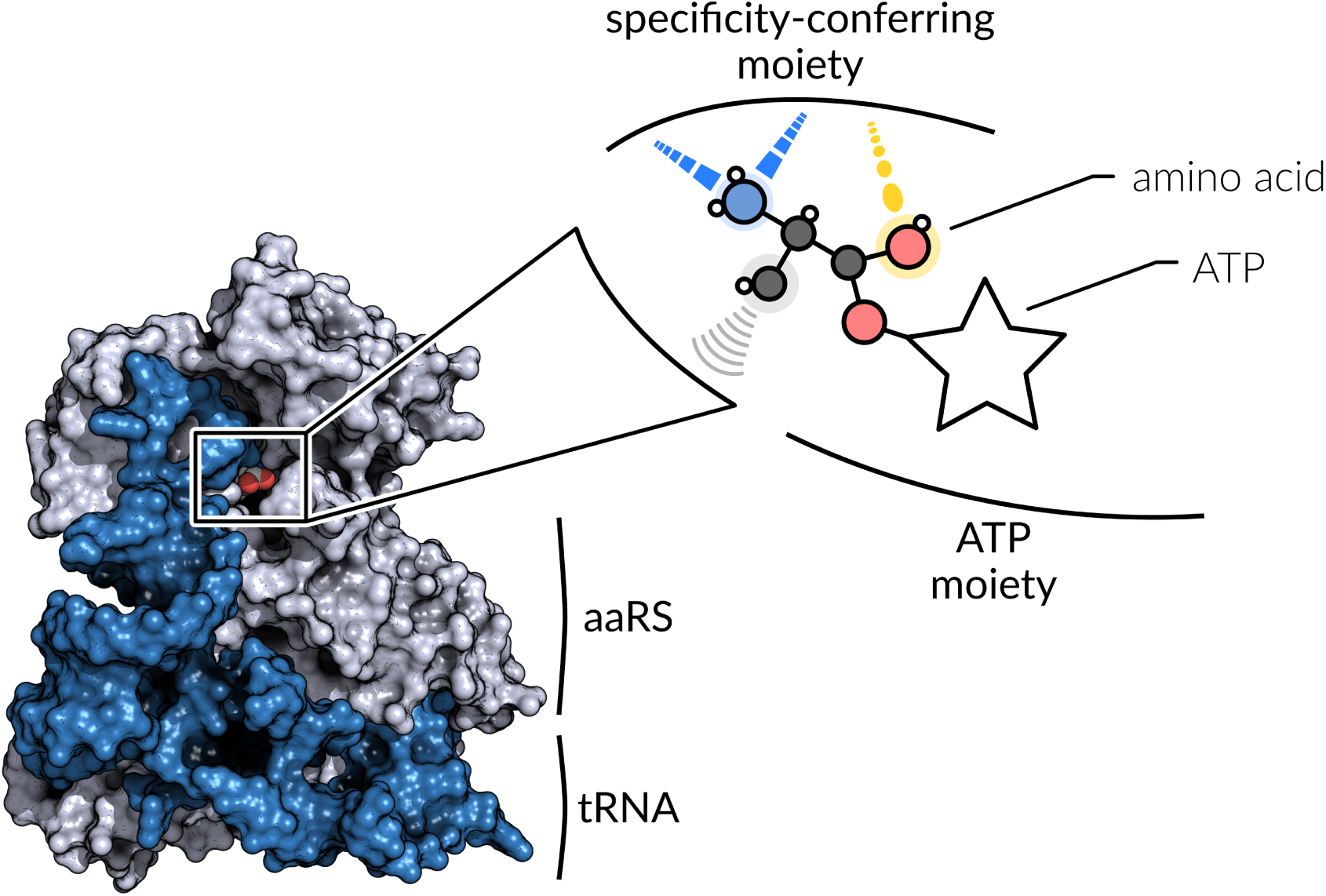
The aaRS tRNA complex and the architecture of its active site. The enzyme catalyzes the covalent attachment of an amino acid to the 3’ end of a tRNA molecule. The binding site itself can be divided into two moieties. While the ATP moiety is responsible for the fixation of ATP, which is consistent within each aaRS class^38^, the specificity-conferring moiety differs between each aaRS and forms highly specific non-covalent interactions with the amino acid ligand.

### Non-Covalent Binding Site Interactions

Non-covalent protein-ligand interactions play an important role for the specific binding of any ligand. These interactions are generally reversible and correspond to an energy of binding between −80 kJ·mol^*−*1^ and −10 kJ·mol^*−*1^, which is less compared to covalent interactions^39^. Several types of non-covalent interactions exist that can add energetic contribution to the binding of a protein ligand-complex. Each type is constrained regarding interaction partners and geometry. Generally, directed hydrogen bonds are considered to be the strongest non-covalent interaction, followed by *π*-cation and *π*-stacking interactions, electrostatic (or salt bridge) interactions, and hydrophobic interactions^40^. Based on experimentally determined three-dimensional structures of protein-ligand complexes, non-covalent interactions can be studied computationally. However, this requires a detailed annotation of non-covalent interaction patterns. In this study, we use the rule-based Protein-Ligand Interaction Profiler (PLIP)^41^ to characterize the amino acid binding in aaRSs.

### Motivation

In our last study we identified two unique ATP binding motifs in Class I and Class II aaRSs^38^, which are by now the minimal description of the two classes. Hence, a detailed study of the amino acid binding site is the logical next step to extend the picture of ligand binding in aaRSs. Protein structures of aaRSs from all kingdoms of life, co-crystallized with their amino acid ligands, are publicly available in the Protein Data Bank (PDB)^42^. Furthermore, there are tools such as PLIP^41^ to characterize and map the interactions of proteins and their ligands. These rich data allow for the investigation of specific characteristics of amino acid recognition in individual aaRS. The overall aim is to contribute to the understanding of how aaRSs realize the correct mapping of the genetic code (Fig. 2) and to provide a compendium of binding site interactions relevant to maintain amino acid specificity. The results shed light on how evolution implemented a specific recognition via the amino acid composition of the binding size, non-covalent interaction patterns, pre- or post-transfer correction mechanisms, and steric effects such as the volume of the binding cavity. Moreover, the overall recognition strategies for Class I and Class II aaRSs differ, suggesting that the existence of the classes allowed the enzymes to cover a broader ligand diversity and thus the gradual incorporation of new amino acids into the genetic code.

**Figure 2.**
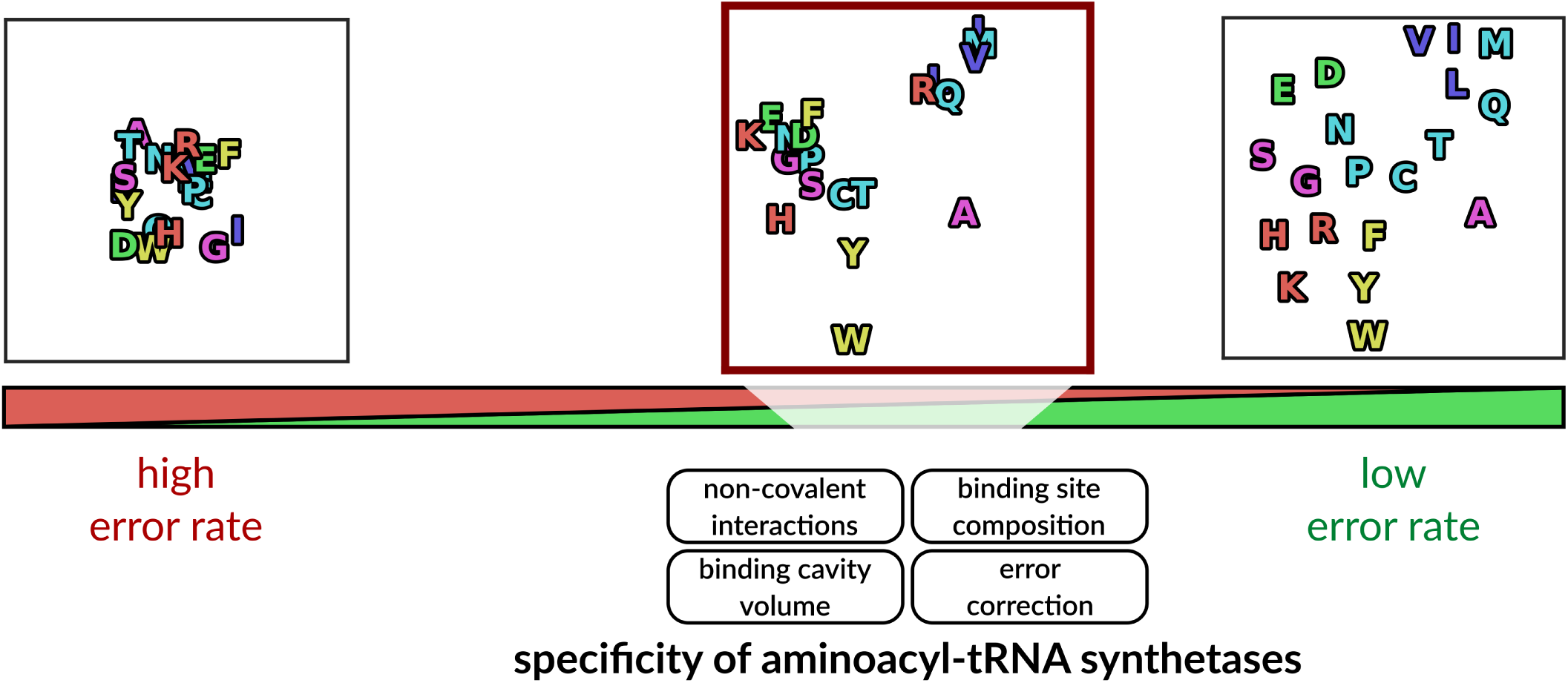
The genetic code relies on the specificity of aminoacyl-tRNA synthetases to ensure the correct mapping of amino acids to their codons. The aaRS enzymes disentangle the recognition space of amino acids to reduce errors during protein synthesis. This study provides a thorough characterization of the mechanisms that drive this specificity. We identified non-covalent interactions in the binding sites of aaRSs, binding site residue composition, error correction mechanisms, and the volume of the binding cavity as key determinants for specific amino acid recognition.

## Results

### Dataset

Based on all available structures in the PDB, 424 (189 Class I, 235 Class II) three-dimensional structures of aaRSs co-crystallized with their corresponding amino acid ligands were analyzed. The selected data covers aaRSs of 56 different species in total, 180 from eukaryotes, 213 from bacteria, and 31 from archaea (SI Appendix Fig. S1). In total, 70 human structures are part of the dataset. Each protein chain that contains a protein-ligand complex of a catalytic aaRS domain was considered. Data was available for each of the 20 aaRSs, plus the non-standard aaRSs pyrrolysyl-tRNA synthetase (PylRS) and phosphoseryl-tRNA synthetase (SepRS). The numbers of protein-ligand complexes available for each aaRS are given in SI Appendix Fig. S2. For twelve aaRSs, protein-ligand complexes were available in both pre-activation and post-activation reaction states, i.e. co-crystallized with either amino acid or aminoacyl ligand (SI Appendix Fig. S3).

### Interaction Features

The frequencies of observed non-covalent binding site interactions in respect of the aaRS class and the type of interaction are shown in Tab. 1. In general, hydrophobic interactions are the most prevalent interactions for Class I aaRSs with a frequency of 44.60% with respect to the total number of interactions, while hydrogen bonds are most frequently observed in Class II aaRSs with 59.23% frequency. Five (hydrogen bonds, hydrophobic interactions, salt bridges, *π*-stacking, and metal complexes) interaction types were observed in aaRSs. No *π*-cation interactions were observed to be involved in amino acid binding. Water-mediated hydrogen bonds were excluded from analyses due to missing data for water molecules for the majority of the crystallographic structures.

### Amino Acid Recognition

The annotation of non-covalent protein-ligand interactions allowed to characterize interaction preferences of each aaRS at the level of individual atoms of their amino acid ligands. This analysis highlights the preferred modes of binding for each of the 22 amino acid ligands. Figure 3 shows the occurring interactions for each aaRS based on the analysis with PLIP. Each interaction is annotated with its occupancy, i.e. the relative frequency of occurrence in respect of the total number of structures for this aaRS. Binding site features are neglected at this point and all interactions are shown with respect to the amino acid ligand.

**Figure 3.**
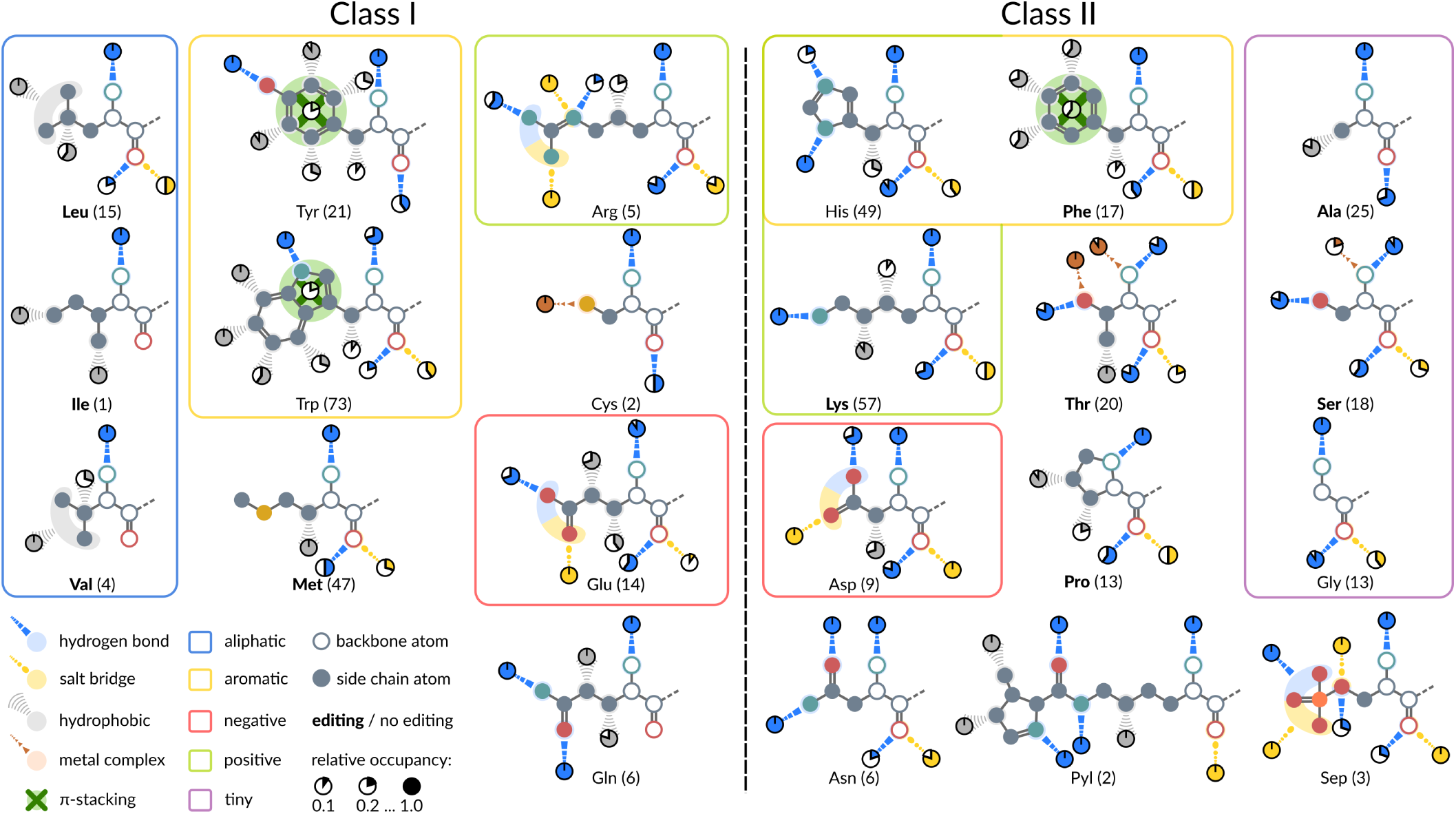
The recognition of individual amino acids by aaRSs mapped to their ligands. The ligands are grouped by physicochemical properties^43^ and aaRS class. Different types of non-covalent protein-ligand interactions were determined with PLIP^41^ and assigned to individual atoms of the ligand using subgraph isomorphism detection^44^. Backbone atoms of the ligand are depicted as circles without filled interior. The relative occupancy of each interaction in respect of the total number of investigated structures (number in parentheses for each aaRS) is given by pie charts. Interactions with an occupancy below 0.1 are neglected. Interactions for which a unique mapping to an individual atom is not possible due to ambiguous isomorphism, e.g. for the side chain of valine, were assigned to multiple atoms. *π*-stacking interactions are shown in dark green and refer to all atoms of the aromatic ring structures in tyrosyl-tRNA synthetases (TyrRSs), tryptophanyl-tRNA synthetases (TrpRSs) and phenylalanine-tRNA synthetases (PheRSs). Some aaRSs prevent the mischarging of their tRNAs via error correction mechanisms (“editing”)^45^. The aaRSs conducting error correction are typeset in bold.

#### Class I

In general, Class I aaRSs interact mainly via hydrogen bonds and hydrophobic interactions with the ligand. The backbone atoms of all Class I ligands feature hydrogen bonding with the primary amine group. The occupancy of this interaction is high throughout all Class I aaRSs, indicating a pivotal role of this interaction for ligand fixation. Additionally, the oxygen atom of the ligand’s carboxyl group is involved in hydrogen bonding except for glutaminyl-tRNA synthetase (GlnRS), isoleucyl-tRNA synthetase (IleRS), and valyl-tRNA synthetase (ValRS). The same atom forms additional salt bridges in leucyl-tRNA synthetase (LeuRS), arginyl-tRNA synthetase (ArgRS), methionyl-tRNA synthetase (MetRS), and glutamyl-tRNA synthetase (GluRS). The side chains of the aliphatic amino acids leucine, isoleucine, and valine are exclusively bound via hydrophobic interactions. ArgRS and GluRS form salt bridges between binding site residues and the charged carboxyl and guanidine groups of the ligand, respectively. Glutamine is bound by GlnRS via conserved hydrogen bonds to the amide group and hydrophobic interactions with beta and delta carbon atoms. The two aromatic amino acids tyrosine and tryptophan are recognized by *π*-stacking interactions and extensive hydrophobic contact networks. Tryptophan is bound preferably from one side of its indole group at positions one, six, and seven. The sulfur atom of the cysteinyl-tRNA synthetase (CysRS) ligand forms a metal complex with a zinc ion in both structures. MetRSs bind their ligand with a highly conserved hydrophobic interaction with the beta carbon atom.

#### Class II

Class II aaRSs consistently interact with the backbone atoms of the ligand via hydrogen bonds and salt bridges. The primary amine group forms hydrogen bonds with high occupancy and is involved in metal complex formation in threonyl-tRNA synthetases (ThrRSs) and seryl-tRNA synthetases (SerRSs). The carboxyl oxygen atoms of the ligands are bound by a combination of hydrogen bonding and electrostatic salt bridge interactions. The overall backbone interaction pattern is highly conserved within Class II aaRSs. Closer investigation revealed that a previously described structural motif of two arginine residues^38^, responsible for ATP fixation, seems to be involved in stabilizing the amino acid carboxyl group with its N-terminal arginine residue. The charged amino acid ligands in histidyl-tRNA synthetase (HisRS) and lysyl-tRNA synthetase (LysRS) form highly conserved hydrogen bonds with the binding site residues. Other specificity-conferring interactions include *π*-stacking interactions and hydrophobic contacts observed for PheRS, metal complex formation for ThrRS and SerRS with zinc, and salt bridges as well as hydrogen bonds for aspartyl-tRNA synthetase (AspRS). The amino acids alanine and proline are bound by alanyl-tRNA synthetases (AlaRSs) and prolyl-tRNA synthetases (ProRSs) via hydrophobic interactions. No specificity-conferring interactions can be described for the smallest amino acid glycine due to absence of a side chain. Hence, glycyl-tRNA synthetase (GlyRS) can only form interactions with the backbone atoms of the ligand. Furthermore, asparaginyl-tRNA synthetases (AsnRSs) mediate highly conserved hydrogen bonds with the amide group of their asparagine ligand. The non-standard amino acid pyrrolysine is bound by PylRS via several hydrogen bonds and hydrophobic interactions with the pyrroline group. SepRSs employ mainly salt bridge interactions to fixate the phosphate group of the phosphoserine ligand.

#### Conserved Interaction Patterns

Class I aaRSs show a strong conservation of hydrogen bonds with the primary amine group of the amino acid ligand with 83.16% of all structures forming this interaction. Interactions with the carboxyl group are less conserved with a frequency of 32.65% for hydrogen bonds and 28.57% for salt bridges, respectively. In this context, the salt bridges with the carboxyl group are a form of extra strong hydrogen bonding^46^. Interaction patterns with the backbone atoms of the amino acid ligand are strikingly consistent within Class II aaRSs. This class forms hydrogen bonds with the primary amine group in 92.15% of all structures. Additionally, hydrogen bonds with the oxygen atom of the carboxyl group occur in 65.70% of all structures and salt bridges with the same atom are formed in 39.26% of all Class II protein-ligand complexes.

#### Similar Recognition Requires Editing Mechanisms

Various aaRSs are known to conduct pre- or post-transfer editing (see the work of Perona and Gruic-Sovulj^45^ for a detailed discussion of editing mechanisms) in order to ensure proper mapping of amino acids to their cognate tRNAs. The similarity of interaction preferences depicted in Fig. 3 suggests that groups of very similar amino acids require editing mechanisms for their correct handling. Especially the three aliphatic amino acids isoleucine, leucine, and valine are bound via unspecific and weak hydrophobic interactions, substantiating the necessity of editing mechanisms observed for their aaRSs^47^ and that substrate hydrophobicity cannot entirely account for specificity^48^. A similar trend can be observed, e.g., for AlaRS^49^ in order to distinguish alanine from serine or glycine.

#### Binding Site Geometry and Cavity Volume

We investigated binding site geometry and cavity volume in order to quantify their potential contribution to amino acid recognition. Known editing mechanisms in aaRSs are focused on the prevention or correction of tRNA mischarging within one aaRS class (intra-class), e.g. the amino acids isoleucine, leucine, and valine belong to Class I. However, GluRSs and AspRSs have a highly similar interaction pattern of hydrogen bonds and salt bridges with the carboxyl group and weak hydrophobic interactions. Both aaRSs do not use editing and are handled by different aaRS classes. In this case, the geometry and size of the binding site can act as an additional layer of selectivity; a mechanism also exploited by ValRS^47, 50^. To quantify the contribution of binding site geometry, seven structures of GluRS and six structures of AspRS were superimposed with respect to their common adenine substructure using the Fit3D^51^ software. As this superimposition can solely be computed for protein-ligand complexes which resemble the post-reaction state, only a subset of the structures was used. The results show that the ligands of GluRSs and AspRSs are oriented towards different sides of a plane defined by their common adenine substructure (Fig. 4A). There is a significant difference (Mann-Whitney U *p<*0.01) in ligand orientation, described by the torsion angle between phosphate and the amino acid substructure of the ligand (Fig. 4B). Class I GluRSs feature a torsion angle of 54.64 *±* 7.12^*°*^, whereas the torsion angle of Class II AspRSs is −65.02 *±* 7.40^*°*^. Furthermore, the volume of the specificity-conferring moiety of the binding site (see Fig. 1) was estimated with the POVME^52^ algorithm. It differs significantly (Mann-Whitney U *p<*0.01) between GluRS (147.00 *±* 22.31 Å^3^) and AspRS (73.34 *±* 17.12 Å^3^). This trend can be observed for all Class I and Class II structures, respectively. An analysis of all representative structures for Class I and Class II aaRSs shows that Class I binding sites are significantly (Mann-Whitney U *p<*0.01) larger on average (Fig. 4C). While Class I binding cavities have a mean volume of 143.40 *±* 39.62 Å^3^, Class II binding sites are on average 90.36 *±* 32.09 Å^3^ in volume.

**Figure 4.**
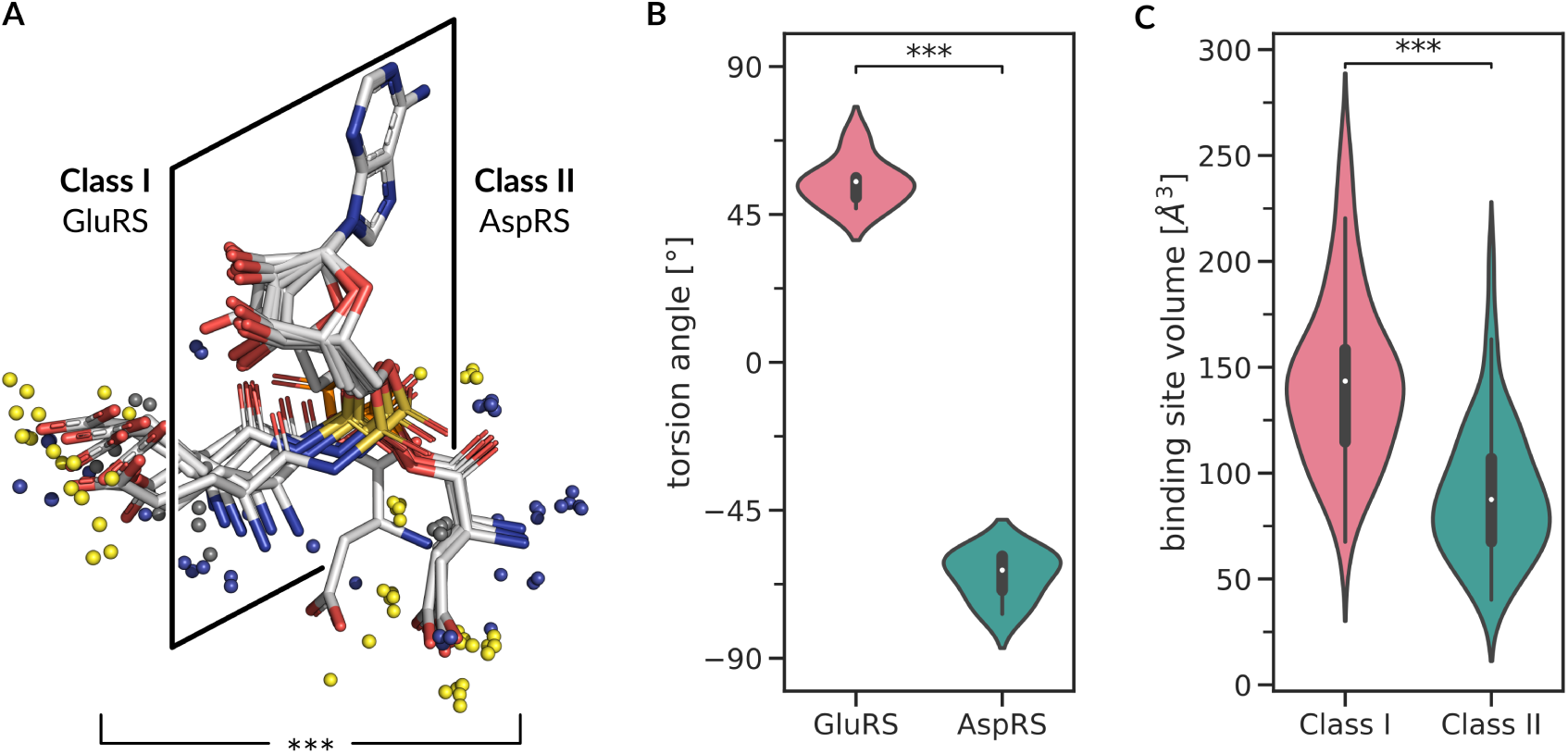
Binding geometry and binding cavity volume analysis. (**A**) Binding geometry of GluRSs and AspRSs. Aminoacyl ligands of Class I GluRSs and Class II AspRSs in post-activation state aligned with Fit3D^51^ with respect to their adenine substructure. The midpoints of non-covalent interactions^41^ with binding site residues are depicted as small spheres. Blue is hydrogen bond, yellow is salt bridge, and gray is hydrophobic interaction. (**B**) Distribution of torsion angles between the phosphate and amino acid substructure of the ligand. The orientation of the ligand in the binding site differs significantly (Mann-Whitney U *p<*0.01) between GluRSs and AspRSs. (**C**) The volume of the specificity-conferring moiety of the binding site, estimated with the POVME algorithm^52^, differs significantly between Class I and Class II aaRSs (Mann-Whitney U *p<*0.01).

### Interaction Patterns of Individual aaRSs

In addition to the investigation of interaction preferences from the ligand point-of-view, the binding sites of each aaRS were analyzed regarding the residues that form interactions with the amino acid ligand. The interactions were mapped to a unified sequence numbering for each aaRS, which is based on multiple sequence alignments (MSAs) (see Methods and Data Availability). Original sequence numbers for each position can be inferred with mapping tables published along with this manuscript (see Data Availability). Figure 5A shows a sequence logo^53^ representation of binding site interactions for AlaRS. Each colored position in the sequence logo represents interactions occurring at this position. Highly conserved interactions can be observed at renumbered position 135. The corresponding hydrogen bond and salt bridge interactions are formed with the backbone atoms of the ligand. On the protein side, this interaction is mediated by a conserved arginine residue that corresponds to the N-terminal residue of the previously described Arginine Tweezers motif^38^. Another prominent interaction is formed by valine at renumbered position 293. This residue interacts with the beta carbon atom of the alanine ligand via hydrophobic interactions. In some structures, this hydrophobic interaction is complemented by an alanine residue at renumbered position 325. Aspartic acid at renumbered position 323 is highly conserved in AlaRSs and seems to be involved in amino acid fixation via hydrogen bonding of the primary amine group. Overall, the specificity-conferring interactions with the small side chain of alanine are hydrophobic contacts. An example for amino acid recognition in AlaRSs is given in Fig. 5B. The structure of bacterial *Escherichia coli* AlaRS forms the whole array of observed interactions. Sequence logos of the remaining aaRSs are given in SI Appendix Fig. S4-24. Based on the interactions between binding site residues and the ligand, a qualitative summary of specificity-conferring mechanisms and key residues was composed (Table 2). Moreover, the ligand size and count of observed interactions was checked for dependence. There is a weak but significant positive correlation between the average number of interacting binding site residues for each aaRS and the number of all non-hydrogen atoms of the amino acid ligand (Pearson *r*=0.32, *p*<0.01). This indicates that the number of formed interactions generally increases with ligand size. However, smaller amino acids do not necessarily have a less complex recognition pattern. ThrRSs, for example, bind their amino acid ligand with on average more than a dozen binding site residues, while ValRSs employ on average five binding site residues. The hydroxyl group of threonine allows for an extended range of non-covalent interactions to be formed with binding site residues compared to valine, where only hydrophobic contacts can be established. Distributions of interacting binding site residues for each aaRS are given in SI Appendix Fig. S25.

**Table 1.** Overview of observed interactions between aaRSs and their amino acid ligands. The most prevalent interactions are hydrophobic interactions for Class I aaRSs and hydrogen bonds for Class II aaRSs (typeset in bold). Relative frequencies in respect of all interactions of the aaRS class are given in parentheses.

|  | interaction type |  |  |  |  |  |
| --- | --- | --- | --- | --- | --- | --- |
| | hydrogen bond | hydrophobic | salt bridge | π-stacking | metal complex | total |
| **Class I** | 468 (37.96%) | **550** (44.60%) | 153 (12.41%) | 59 (4.79%) | 3 (0.24%) | 1233 |
| **Class II** | **856** (59.23%) | 193 (13.36%) | 202 (13.98%) | 144 (9.97%) | 50 (3.46%) | 1445 |

**Table 2.**
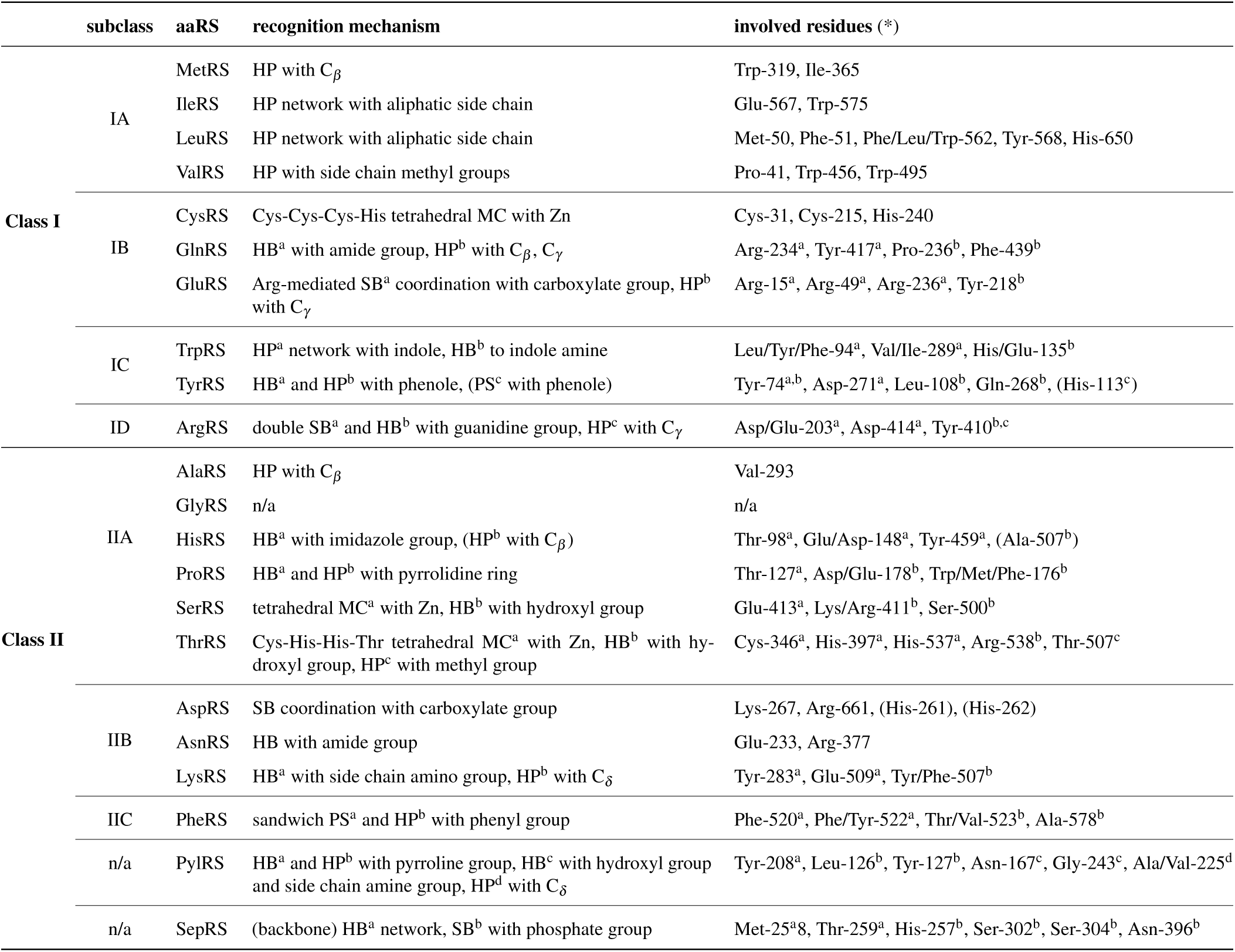
Overview of specificity-conferring recognition mechanisms for all aaRSs grouped by aaRS class and subclass^15^. Only interactions with side chain atoms of the amino acid ligand were included in this summary. HB is hydrogen bond, SB is salt bridge, HP is hydrophobic, MC is metal complex, and PS is *π*-stacking interaction. Correspondences between interactions and residues are indicated by superscript letters. Entries in parentheses were only observed in certain structures and are no general pattern. (*) Residue numbers are given according to the respective MSA (see Methods). Original residue numbers can be inferred with tables published along with this manuscript (see Data Availability).

**Figure 5.**
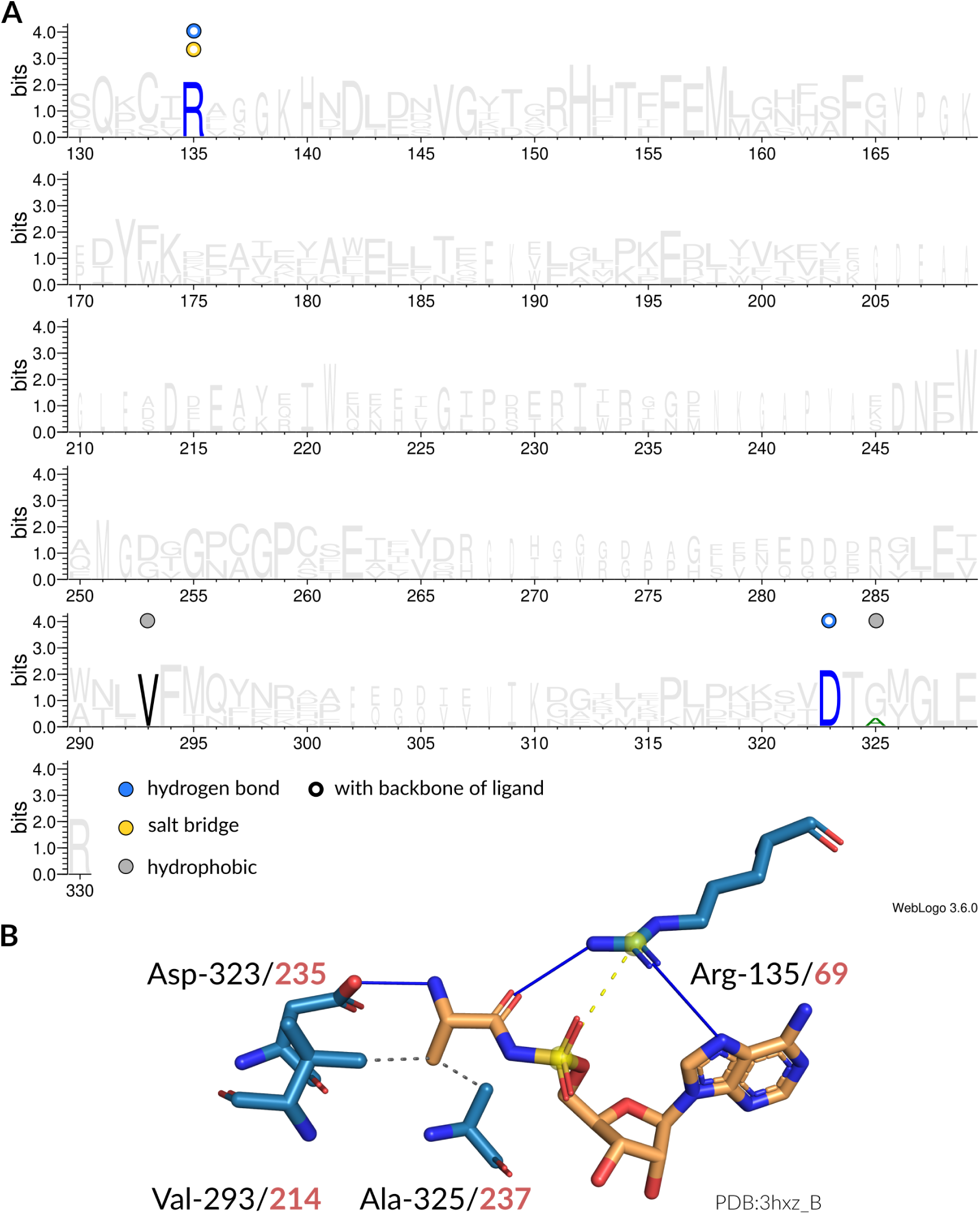
Interaction patterns of AlaRS. (**A**) Sequence logo^53^ of representative sequences for AlaRSs. Non-covalent interactions with the amino acid ligand occurring at certain positions are indicated by colored circles. Filled circles are interactions with the side chain atoms, while hollow circles are interactions with any of the backbone atoms of the amino acid ligand. Blue is hydrogen bond, yellow is salt bridge, gray is hydrophobic interaction. (**B**) Depiction of interactions in the binding site (blue stick model) of an AlaRS from *Escherichia coli* (PDB:3hxz chain A) with its ligand (orange stick model). Here, hydrogen bonds (solid blue lines) and hydrophobic interactions (dashed gray lines) are established. The sequence positions of the interacting residues are given in accordance to the MSA (black) as well as the original structure (red). Figure created with PyMol^54^. Double bonds are indicated by parallel line segments, aromatic bonds by circular dashed lines.

### Quantitative Comparison of Ligand Recognition

To allow for a quantitative analysis and comparison of ligand recognition between several aaRSs, interaction and binding site features were represented as binary vectors, so-called interaction fingerprints (see Methods). Based on these fingerprints, the Jaccard distance was computed for each pair of structures to represent the dissimilarity in ligand recognition. Subsequently, the Uniform Manifold Approximation and Projection for Dimension Reduction (UMAP) algorithm^55^ was used for dimensionality reduction and embedding of the high-dimensional fingerprints into two dimensions for visualization. This embedding is considered to be the *recognition space* of aaRSs. Figure 6A shows the embedding results for all aaRSs in the dataset colored according to the aaRS classes. A Principal Component Analysis (PCA) of the same data is given in SI Appendix Fig. S26. For each aaRS the average position of all data points in the embedding space was calculated and is shown as one-letter code label. Fig. 6B shows the same data colored according to the physicochemical properties of the amino acid ligand, i.e. positive (lysine, arginine, and histidine), aromatic (phenylalanine, tyrosine, and tryptophan), negative (aspartic acid and glutamic acid), polar (asparagine, cysteine, glutamine, proline, serine, and threonine), and unpolar (glycine, alanine, isoleucine, leucine, methionine, and valine).

**Figure 6.**
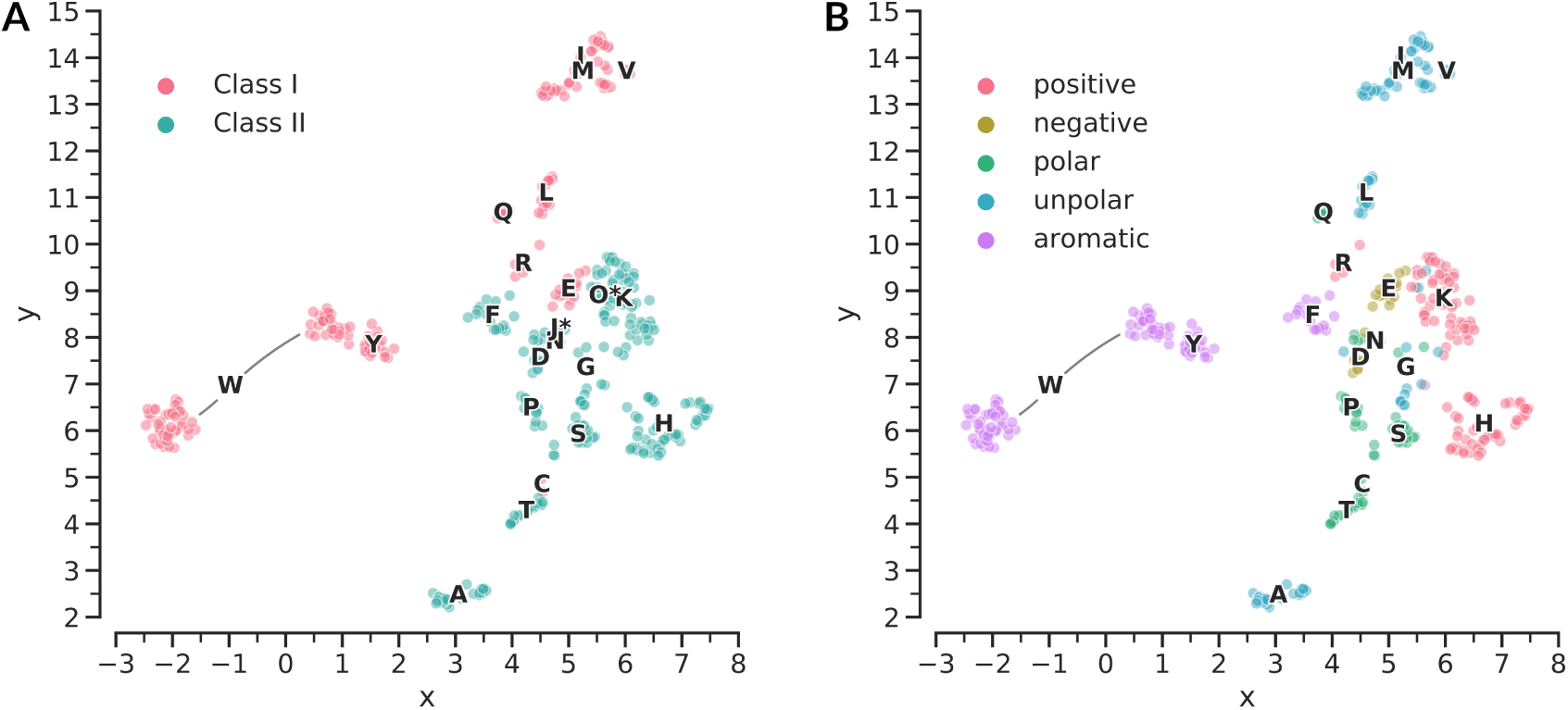
Recognition space analysis of all aaRSs. (**A**) Embedding^55^ space of interaction fingerprints for all aaRS structures in the dataset. Scaling is in arbitrary units. The data points are colored according to the aaRS class. One letter code labels are given for each aaRS based on the averaged coordinates in the embedding space. An asterisk indicates the non-standard amino acids phosphoserine (J*) and pyrrolysine (O*). (**B**) Embedding space of interaction fingerprints for all aaRS structures in the dataset except phosphoserine and pyrrolysine. Scaling is in arbitrary units. One-letter codes of amino acid ligands are used to identify each aaRS. Every data point represents an individual protein-ligand complex. The color of the data points encodes the physicochemical properties^43^ of the ligand.

#### Class I

In terms of amino acid binding both aaRS classes seem to employ different overall mechanism; they separate almost perfectly in the embedding space. Especially aromatic amino acid recognition in Class I TrpRSs and TyrRSs is distinct from Class II aaRSs and forms two outgroups in the embedding space. Remarkably, two different recognition mechanisms exist for TrpRSs, indicated by two clusters approximately at positions (−2.0,6.0) and (1.0,8.5) of the embedding space, respectively. The cluster at position (−2.0,6.0) is formed by structures from bacteria and archaea, while the cluster at position (1.0,8.5) is formed by eukaryotes and archaea and is in proximity to TyrRSs. Closer investigation of two representatives from these clusters shows two distinct forms of amino acid recognition for TrpRSs. Human aaRSs employ a tyrosine residue in order to bind the amine group of the indole ring, while prokaryotes employ different residues (SI Appendix Fig. S27). The Class I aaRSs that are closest to Class II are GluRSs and CysRSs. A cluster of high density is formed by Class I IleRS, MetRS, and ValRS, which handle aliphatic amino acids. This indicates closely related recognition mechanisms and difficult discrimination between these amino acids.

#### Class II

For Class II aaRSs the recognition space is less structured. Nonetheless, clusters are formed that coincide with individual Class II aaRSs, e.g. a distinct recognition mechanism in AlaRSs. The aaRSs handling the small and polar amino acids threonine, serine, and proline are closely neighbored in the embedding space. Recognition of GlyRSs seems to be diverse; GlyRSs are not grouped in the embedding space. However, the recognition of glycine, which has no side chain, is limited by definition and thus the fingerprinting approach might fail to capture subtle recognition features. AspRSs and AsnRSs are located next to each other in the embedding space. Their recognition mechanisms seem to be very similar as the only difference between these two amino acids is the carboxylate and amide group, respectively.

#### Mechanisms That Drive Specificity

In order to quantify the influence of different aspects of binding site evolution on amino acid recognition by aaRSs, different interaction fingerprint designs were compared against each other. Each design includes varying levels of information and combinations thereof: the sequence composition of the enzyme’s binding site (Seq), non-covalent interactions formed between side chains of the enzyme’s binding site and the amino acid ligand (Int), whether pre- or post-transfer correction (i.e. “editing”) is conducted (Ed), and the overall volume of the enzyme’s binding cavity (Vol). To assess the segregation power of each fingerprint variant, the mean silhouette coefficient^56^, a quantification for the error in clustering methods, over all data points was calculated. This score allows to assess to which extent the recognition of one aaRS differs from other aaRSs and how similar it is within its own group. Perfect discrimination between all amino acids would give a value close to one, while a totally random assignment corresponds to a value of zero. Negative values indicate that the recognition of a different aaRS is rated to be more similar than the recognition of the same aaRS. Figure 7 shows the results of this comparison. When using fingerprints describing the sequence composition of the enzyme’s binding site (Seq_sim_), the mean silhouette coefficient over all samples is −0.0510, which indicates many overlapping data points and unspecific recognition. By including non-covalent interactions (Seq, Int) the value increases to 0.1361. If pre- or post-transfer correction mechanisms are considered (Seq, Int, Ed), the silhouette coefficient improves further to 0.2731. Adding information about the binding cavity volume (Seq, Int, Ed, Vol) slightly increases the quality of the embedding to 0.2757. The silhouette coefficients for error correction and volume-based fingerprints were calculated as baseline comparison. If only pre- or post-transfer correction mechanisms (Ed) are considered the mean silhouette coefficient amounts to −0.3027. For binding cavity volume (Vol) the mean silhouette coefficient is −0.4682.

**Figure 7.**
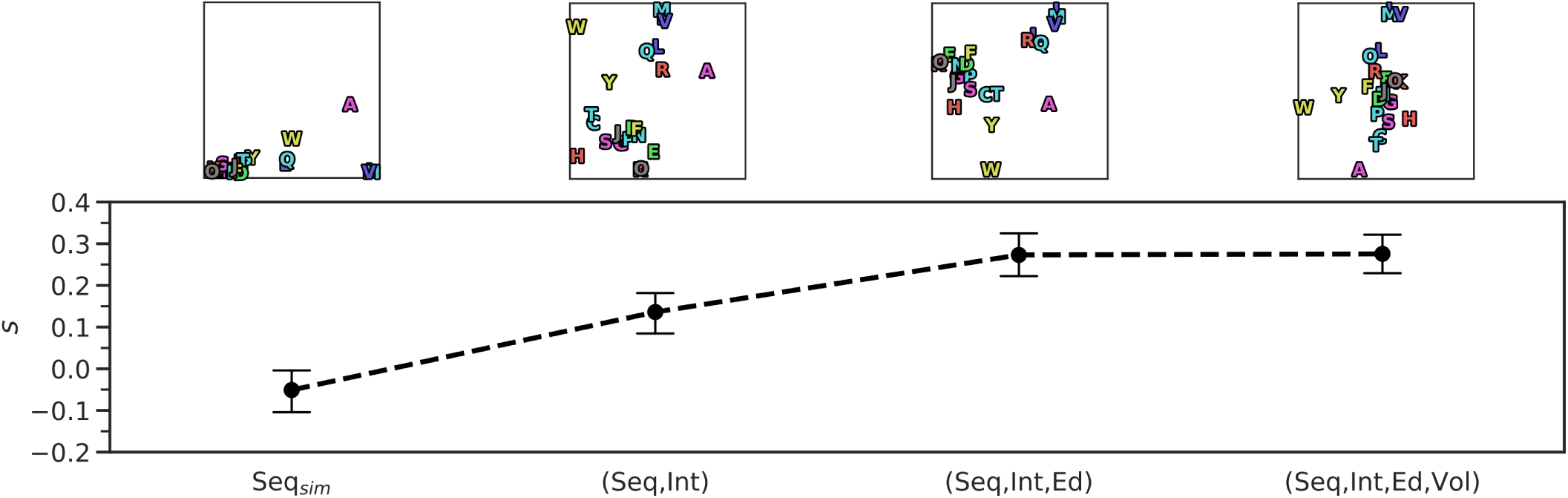
Comparison of different fingerprint designs that include the sequence composition of the enzyme’s binding site (Seq), non-covalent interactions formed between side chains of the enzyme’s binding site and the amino acid ligand (Int), pre- or post-transfer correction (i.e. “editing”) mechanisms (Ed), and volume of the enzyme’s binding cavity (Vol). Simple sequence-based fingerprints (Seq_sim_) are a 20-dimensional representation of binding site composition. The line plot shows the silhouette coefficient^56^ for each embedding. Points represent mean values, error bars are calculated based on all silhouette coefficients for each data point.

#### Relation to Physicochemical Properties of the Ligands

In order to investigate whether the fingerprinting approach is a simple encoding of the physicochemical properties of the amino acids, the results were related to experimentally determined phase transfer free energies for the side chains of amino acids from water (Δ*G*_*w>c*_) and vapor (Δ*G*_*w>c*_) to cyclohexane^3, 57^. These energies are descriptors for the size and polarity of amino acid side chains and underlie both, the rules of protein folding and the genetic code^58^. The Spearman’s rank correlation between pairwise distances for each aaRS in the recognition space and physicochemical property space is weak with *ρ*=0.2564 and *p*<0.01 (see SI Appendix Fig. S28). This indicates that the fingerprinting approach used in this study is a true high-dimensional representation of the complex binding mechanisms of amino acid recognition in aaRSs. This assumption is supported by a PCA (SI Appendix Fig. S26) of the fingerprint data, where the first two principal components account for only 9.24% and 8.44% of the covered variance, respectively.

## Discussion

The correct recognition of individual amino acids is a key determinant for evolutionary fitness of aaRSs and considered to be one of the major determinants for the closure of the genetic code^10^. The results of this study emphasize the multitude of mechanisms that lead to the identification of the correct amino acid ligand in the binding sites of aaRSs. Based on available protein structure data, a thorough characterization of binding site features and interaction patterns allowed to pinpoint the most important drivers for the correct mapping of the genetic code. The main findings of this analysis can be summarized as follows: (i) Class I and Class II aaRSs employ different overall strategies for amino acid recognition. (ii) Interaction patterns and binding site composition are the most important drivers to mediate specificity. However, very similar amino acids require additional selectivity through steric effects or editing mechanisms. (iii) The analysis of interaction fingerprints suggests that error-free recognition is a delicate task demanding a complex interplay between binding site composition, interaction patterns, editing mechanisms, and steric effects. The results point towards a gradual diversification of amino acid recognition and, hence, a gradual extension of the genetic code.

### Class Duality Extends Possibilities

The aaRS class duality allowed to broaden the amino acid recognition space significantly. In general, the recognition of amino acids with low side chain complexity seems to be complemented by allosteric interactions and cannot be exclusively implemented by configuring side chains. Although the volumes of Class I and Class II binding sites differ significantly, they are probably not the major determinants for amino acid selectivity. In general, Class I aaRSs handle most of the hydrophobic and larger amino acids^3^ and thus the binding site volume of Class I aaRSs is expected to match the volumes of their larger ligands. Nonetheless, binding site volume and geometry may act as additional layers of selectivity. An example are the two negatively charged amino acids glutamic acid and aspartic acid, handled by a Class I and Class II aaRS, respectively. In this case, overall interactions are highly similar but binding geometry and binding site volume is significantly different. Both ligands are attacked from the opposite side^59^ as highlighted by significantly different conformations (Fig. 4B). There is evidence that both amino acids were among the first to exist in the prebiotic context^60–65^. It is conceivable that the discrimination between glutamic and aspartic acid was based on superordinate secondary structures elements and size selectivity rather than on specific side chain interactions^66^. The recent identification of a protein folding motif^67^ strengthens this assumption. This is further supported by the observation that ancient proteins, based on a limited set of amino acids, were still capable to exhibit secondary structures^65, 68, 69^. One can only speculate whether a simultaneous emergence of two different aaRS classes and secondary structure formation allowed to incorporate these early – but highly similar – amino acids into the genetic code. According to the biochemical pathway hypothesis^60^, GluRS and AspRS might have been the first Class I and Class II representatives, with other aaRSs evolving from them^60, 70, 71^. However, the decreased usage of aspartic acid and the enrichment of glutamic acid in modern species, compared to the LUCA, points towards a different direction^72^. According to these usage frequencies, aspartic acid was incorporated into the genetic code prior to glutamic acid. This temporal order was equally concluded by the evaluation of various criteria to derive a consensus order of amino acid appearance^73^.

### Glutamine and Asparagine Followed Glutamic Acid and Aspartic Acid

Glutamine and asparagine are chemically closely related to glutamic and aspartic acid, respectively. It is likely that GlnRSs^6^ and AsnRSs^7^ mutually co-evolved from GluRSs and AspRSs through early gene duplication^15^. Although the ligands of GluRS and GlnRS are rather similar, interaction patterns and binding site compositions differ between these two enzymes. These differences coincide with the analysis of the recognition space (Fig. 6), where GluRSs and GlnRSs are not neighbored in the embedding. Hence, they evolved to distinguish between these amino acids without editing mechanisms^74^ or the exploitation of the negative charge of glutamic acid^75, 76^. The discrimination of glutamine and glutamic acid by GlnRS cannot be attributed entirely to the composition of the binding site; changing the specificity from glutamine and glutamic acid could not be achieved by mutating only first order binding site residues^77^. This emphasizes the role of subtler interactions and allosteric effects within the catalytic domain as it was shown to be the case for TrpRS^78^. In contrast to the observed differences between GlnRS and GluRS, AspRS and AsnRS are directly neighbored in the embedding space and share a greater similarity in their recognition mechanism. However, as for GlnRS and GluRS, the discrimination between aspartic acid and asparagine is not entirely driven by specific interactions with binding site residues. Correct recognition depends on a water molecule that forms a water-assisted hydrogen bonding network in the active site of AsnRS^79^. The vicinity in the recognition space might be due to the limitation of interaction data, for which co-crystallized water molecules were not available for the majority of the structures and thus not considered during analysis. We conclude that for both Class I GlnRS/GluRS and Class II AsnRS/AspRS, the role of allosteric effects and other subtle interactions should not be underestimated.

### Distinct Recognition of Arginine and Lysine

Another interesting example are the two positively charged amino acids lysine and arginine. Interaction data suggests two unrelated ways to achieve ligand recognition in Class II LysRSs and Class I ArgRS, i.e. the two enzymes are well separated in the embedding space. The poor editing capabilities for LysRS regarding arginine^80^ might have required a good separation of the two recognition mechanisms. Even if a relation of ArgRSs to aaRSs of hydrophobic amino acids was proposed^81^, a separate subclass grouping for ArgRSs^15^ seems to be reasonable and is in accordance with the observed data; the recognition mechanism differs substantially from the hydrophobic amino acids. Furthermore, based on the consensus of all analyzed ArgRS structures, the characteristic Class I HIGH motif^4^ seems to play an important role for stabilization of the arginine ligand in pre-activation state (see SI Appendix Fig. S4). For both histidine residues of the HIGH motif highly conserved salt bridges are observed that bind to the carboxyl group of the ligand.

### Glycine Recognition is not Interaction-Driven

Based on interaction data, the recognition of the smallest amino acid glycine seems to be rather unspecific; a large spread in the embedding space can be observed for individual protein-ligand complexes of GlyRS. This is to be expected as GlyRS is known to maintain its specificity not due to interactions with glycine – it has no side chain to interact with – but rather due to active site geometry that blocks larger amino acids^10, 82^.

### Alanine Recognition is Crucial

Alanine is the second smallest amino acid with only a single heavy side chain atom. The idiosyncratic architecture of AlaRS is different from other Class II aaRSs^83^. Still, the confusion with glycine and serine^49^, or non-proteinogenic amino acids^8^, poses a challenge for correct recognition of alanine and a loss of specificity is associated with severe disease outcomes^84^. The recognition mechanism in AlaRSs seems to differ substantially from other Class II aaRSs (see Fig. 6), indicating evolutionary endeavor to develop a unique recognition mechanism.

### Discrimination of Hydrophobic Amino Acids Requires Editing

The hydrophobic amino acids isoleucine, leucine, valine, and methionine likely entered the genetic code at the same time^20, 61, 81^. The highly similar interaction patterns for IleRS, ValRS, and MetRS substantiate this assumption. Due to their difficult discrimination, editing functionality is key^5, 50, 74, 85, 86^ for these aaRSs.

### Tryptophan Recognition Suggests Late Addition to the Genetic Code

The emergence of TrpRSs and TyrRSs is considered to have happened at a later stage of evolution. The two aaRSs are likely to be of common origin^37, 87^ and constitute their own subclass, which is supported by sequence and structure studies^15, 18, 19, 88, 89^. PheRS supposedly evolved from the same precursor as TrpRS and TyrRS^21^. In general, TrpRSs and TyrRSs separate well from other aaRSs in the recognition space, which is likely due to the unique utilization of *π*-stacking interactions with binding site residues. Beside specific interactions in the binding site, allosteric effects and interdomain cooperativity^90, 91^ are drivers for TrpRS specificity. Furthermore, mutations in the dimerization interface of TrpRSs were shown to reduce specificity^78^. Remarkably, two distinct ways of recognition are apparent for TrpRSs in bacteria and eukaryotes. These differences support the previous described separation of eukaryotic TrpRSs and TyrRSs from their prokaryotic counterparts^92^ and late addition of these amino acids to the genetic code^93^. However, structures from archaea do not follow this pattern and feature both recognition variants.

## Methods

### Data Acquisition

The dataset from our last study^38^ served as the basis for all analysis. As all structures in the dataset are annotated with ligand information, only entries containing ligands relevant for amino acid recognition were considered, i.e. they bind to the specificity-conferring moiety of the binding site (see Fig. 1). Every protein chain of the entry was considered that: (i) comprises a catalytic aaRS domain, (ii) contains a co-crystallized specificity-relevant ligand in the active site, and (iii) the ligand must contain an amino acid substructure. Filtering of the data resulted in 189 (235) structures for Class I (Class II) aaRSs that contain ligands with relevance for specificity. The number of structures in respect of the pre- or post-activation state of the catalyzed reaction is shown in SI Appendix Fig. S3. Furthermore, sequences of the dataset entries were clustered using single-linkage clustering with a sequence identity cutoff of 95% according to a global Needleman-Wunsch^94^ alignment with BLOSUM62 substitution matrix computed with BioJava^95^. Representative chains for each cluster were selected, preferring wild type and high-quality structures. In total, 47 (54) protein chains were selected to be representatives for Class I (Class II) aaRSs. The dataset covers structures of all known aaRSs from species across all kingdoms of life (SI Appendix Fig. S1).

### Mapping of Sequence Positions

Amino acid sequences were derived from the set of representative structures of the respective aaRS. To allow a unified mapping of sequence positions, an MSA was computed for each aaRS using the T-Coffee^96^ Expresso pipeline. The quality of each MSA in the specificity-conferring region of the binding site was assessed regarding the correct mapping of the Backbone Brackets and Arginine Tweezers structural motifs^38^, and the conservation of the respective sequence signature motifs^4, 22^. All MSAs preserved the considered regions and passed the quality checks. The sequence positions for each aaRS were then unified according to the resulting MSA in order to investigate conserved interaction patterns. For this purpose the custom script “MSA PDB Renumber”, available under open-source license (MIT) at github.com/vjhaupt, was used.

### Annotation of Non-Covalent Protein-Ligand Interactions

Non-covalent protein-ligand interactions were annotated for all entries in the dataset that contained a valid ligand using PLIP v1.3.3^41^ with default parameters.

### Determination of Interactions Relevant for Specificity

Only interactions formed between the amino acid substructure of the ligand and binding site residues were considered for analysis. For this purpose subgraph isomorphism detection with the RI algorithm^44^ was applied. The RI implementation of the SiNGA framework v0.5.0^97^ was used. Each amino acid scaffold was represented by a graph created from the amino acid’s SMILES string taken from PubChem^98^. The full amino acid graph was modified using MolView v2.4 (available at molview.org) in order to remove the terminal hydroxyl group, which is cleaved during the enzymatic reaction and must thus be ignored for subgraph matching. For each dataset entry that contained a valid ligand, the corresponding amino acid graph was matched against the ligand in order to identify the atoms involved in the formation of specificity-conferring interactions. A depiction of the workflow to determine specificity-conferring interactions is given in Fig. 8.

**Figure 8.**
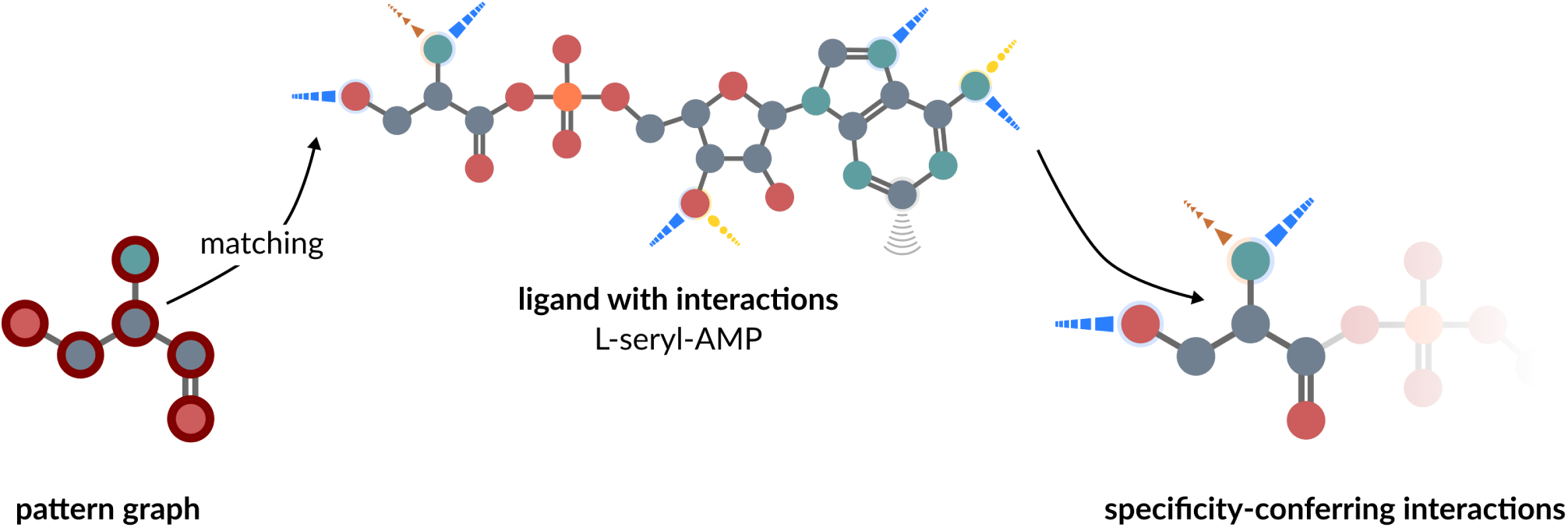
The identification of specificity-conferring interactions in SerRS. For each aaRS a pattern graph is used to map interactions. This patterns graph resembles the amino acid without its terminal hydroxyl group and is matched against the full ligand with annotated interactions using subgraph isomorphism detection^44^. The interactions formed between matched atoms and binding site residues are considered to be specificity-conferring interactions.

### Generation of Interaction Fingerprints

To allow for a quantitative comparison of recognition mechanisms, each protein-ligand complex was represented by a structure-invariant binary interaction fingerprint (see for example the paper of Salentin *et al*.^40^ about the idea of interaction fingerprinting). Different fingerprint designs were chosen for comparison: a simple 20-dimensional fingerprint on binding site composition and a 500-dimensional fingerprint based on binding site composition and interaction information. The latter was further enriched with editing and binding site volume information.

#### Simple Binding Site Based Fingerprints

Binary and structure-invariant fingerprints that represent binding site compositions (used as baseline for the comparison of different fingerprint designs, Fig. 7) were constructed as follows. Each residue predicted to be in contact with any specificity-relevant atom of the ligand was considered for fingerprint generation. A 20-dimensional binary vector was used to represent the occurrence of individual residue types in the binding site. For each of the interacting residues the corresponding bit was set to active. Hence, multiple occurrences of the same residue type were not taken into account.

#### Binding Site and Interaction-Based Fingerprints

Single three-dimensional vectors of non-covalent interactions were encoded into a binary vector by considering the type of interaction, the interacting group in the ligand and the interacting amino acid residue. One such feature could be a hydrogen bond between an oxygen atom in the ligand and tyrosine in the protein. Each of these features is hashed to a number between 1 and 500 so that the resulting fingerprint has 500 bits.

#### Encoding of Editing Mechanisms and Binding Site Volume

Information about the editing mechanisms performed by some aaRSs were taken from the paper of Perona and Gruic-Sovulj^45^ and encoded by appending a 22-dimensional bit vector to the 500-dimensional fingerprint. Each active bit represents a ligand against which editing is performed, e.g. for structures of ThrRS the bit for serine is set. In addition to editing information the binding site volume, estimated with the POVME^52^ algorithm, was encoded. Twelve bins were created that represent binding site volumes ranging from 30-270 Å^3^ in steps of 20 Å^3^. For example, if a structure has a binding site volume of 45 Å^3^ the first bit was set to active. For a binding site volume of, e.g., 52 Å^3^ the second bit was set to active and so on. The fingerprints were concatenated to contain the binding site and interaction features (500 bits), editing mechanisms (22 bits), and binding site volume (12 bits). The final fingerprint has a size of 534 bits.

### Embedding of Interaction Fingerprints

To allow for a quantitative comparison of the interactions between individual aaRSs, the high-dimensional interaction fingerprints were embedded using UMAP version 0.3.2^55^. The parameters for all embeddings given in this manuscript were set as follows: `n_neighbors = 60`, `min_dist = 0.1`, `n_components = 2`. The Jaccard distance was used to describe the dissimilarity between two fingerprints *a* and *b*:

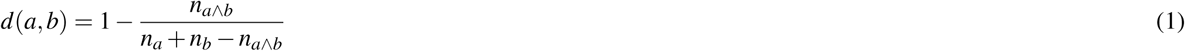

with *n*_*a∧b*_ being the count of active bits common between fingerprints *a* and *b, n*_*a*_ the number of active bits in fingerprint *a*, and *n*_*b*_ the number of active bits in fingerprint *b*. This distance metric was used as input for UMAP.

## Data Availability

All accompanying data is made publicly available under 10.5281/zenodo.3598250. This repository contains the MSA files of representative structures for each aaRS that were used for consistent renumbering as well as Excel tables to infer original sequence positions from renumbered positions for each aaRS. Rows are renumbered positions, columns are sequence positions of individual structures.

## Author contributions statement

F.K. and S.K. prepared and analyzed data. F.K., S.K, and S.S. wrote the manuscript. S.S. designed and computed interaction fingerprints, V.J.H. implemented renumbering of structures, C.L. and S.B. supported data curation, formal analysis, and conceptualization. D.L. and M.S supervised the project. All authors read and approved the manuscript.

## Additional information

The authors declare no conflict of interest.

## Supplementary Information

**Fig. S1.**
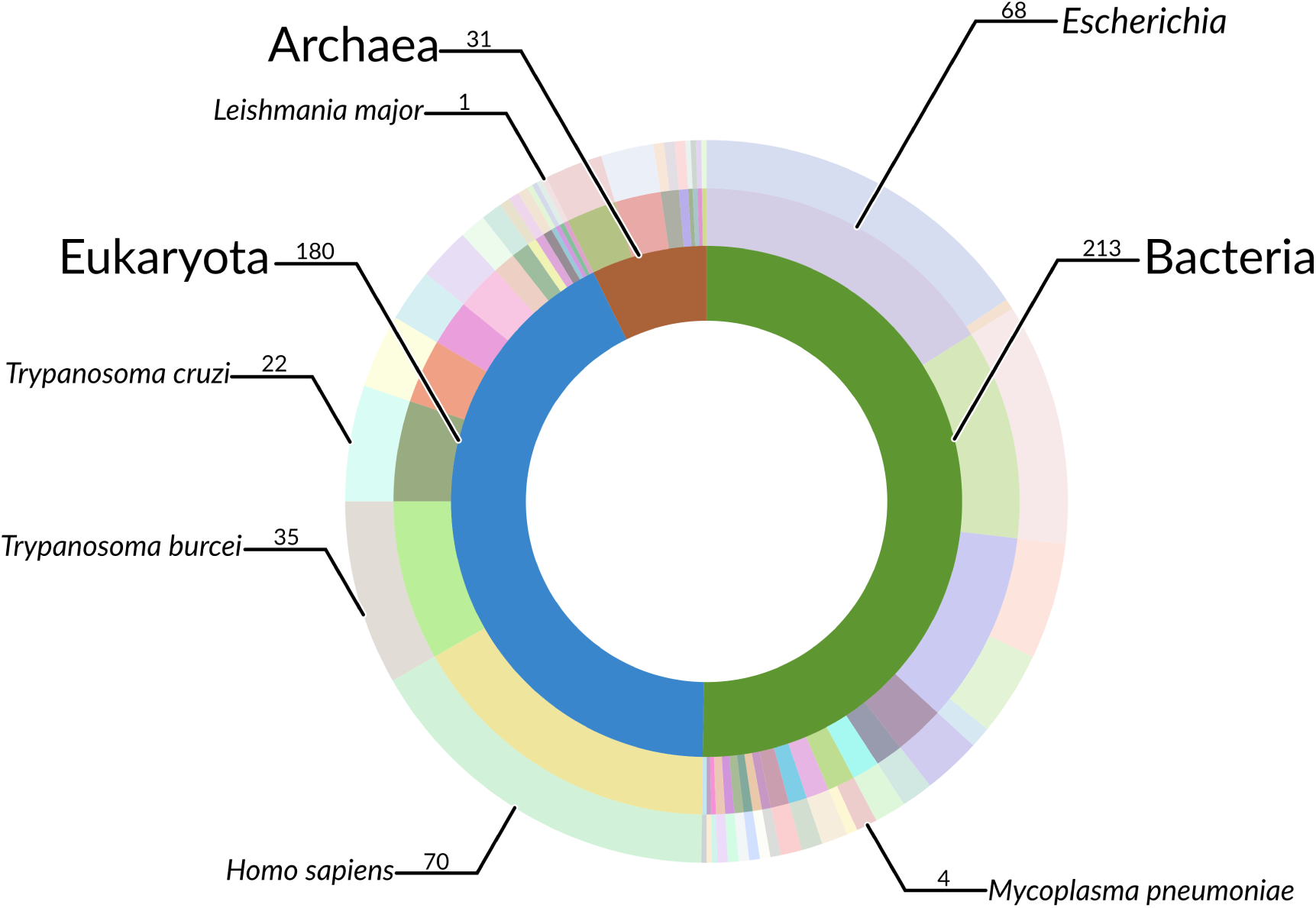
Sunburst diagram of protein chains containing a catalytic Aminoacyl-tRNA synthetase (aaRS) domain co-crystallized with their amino acid ligand in respect to source species. The dataset (1) covers all three superkingdoms, contains human and structures of pathogenic species.

**Fig. S2.**
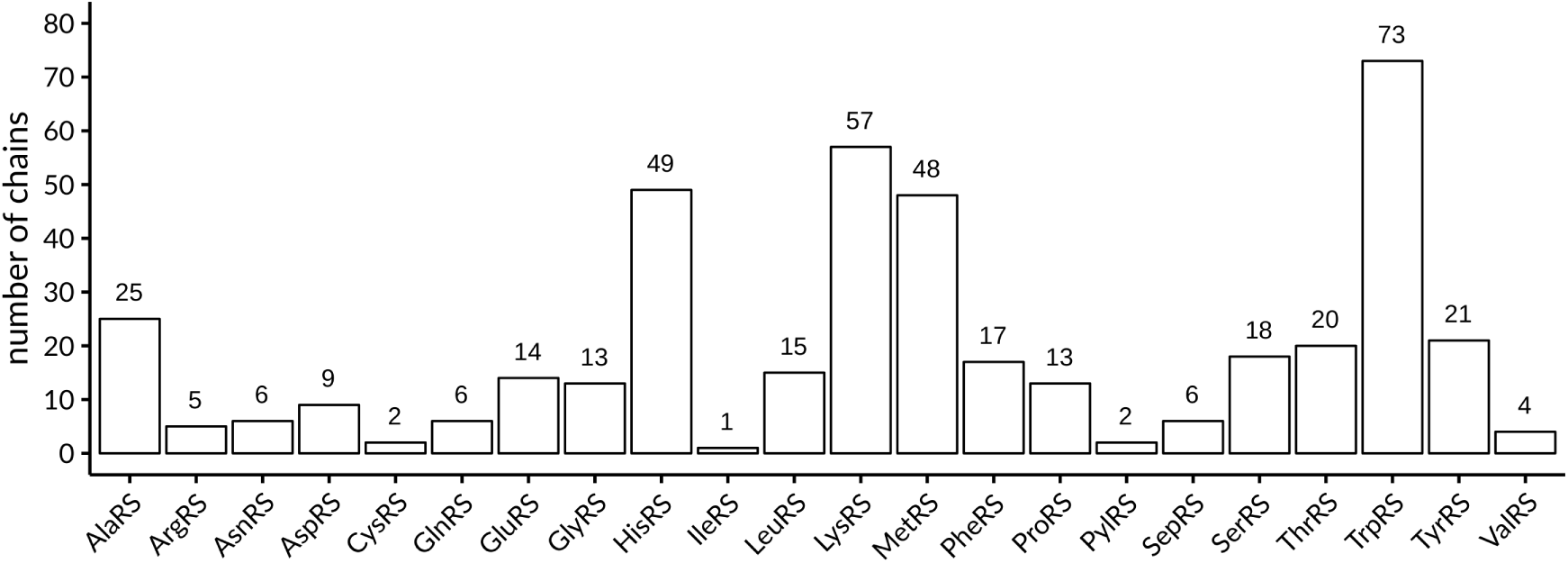
The number of protein chains containing a catalytic aaRS domain for each of the 22 aaRSs. The dataset(1) used in this study contains structures for all aaRSs.

**Fig. S3.**
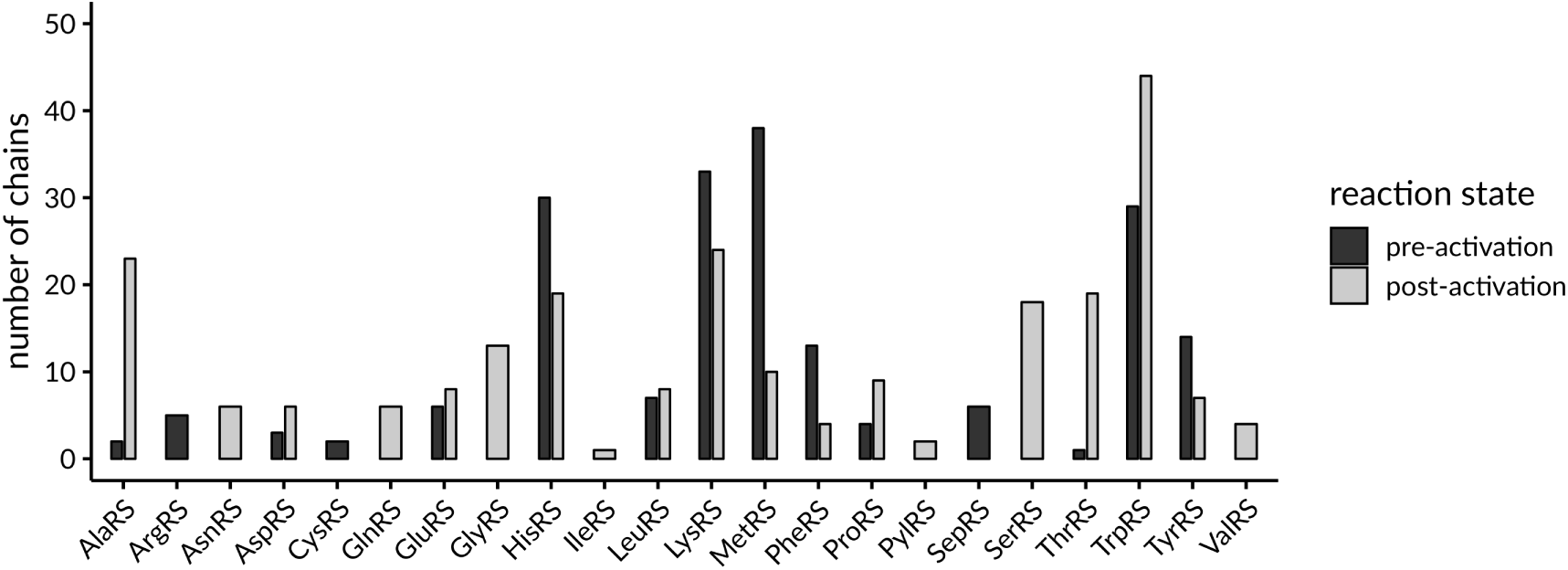
The number of protein chains containing either an amino acid ligand (pre-activation) or an aminoacyl ligand (post-activation). For twelve aaRSs data was available for both reaction states.

**Fig. S4.**
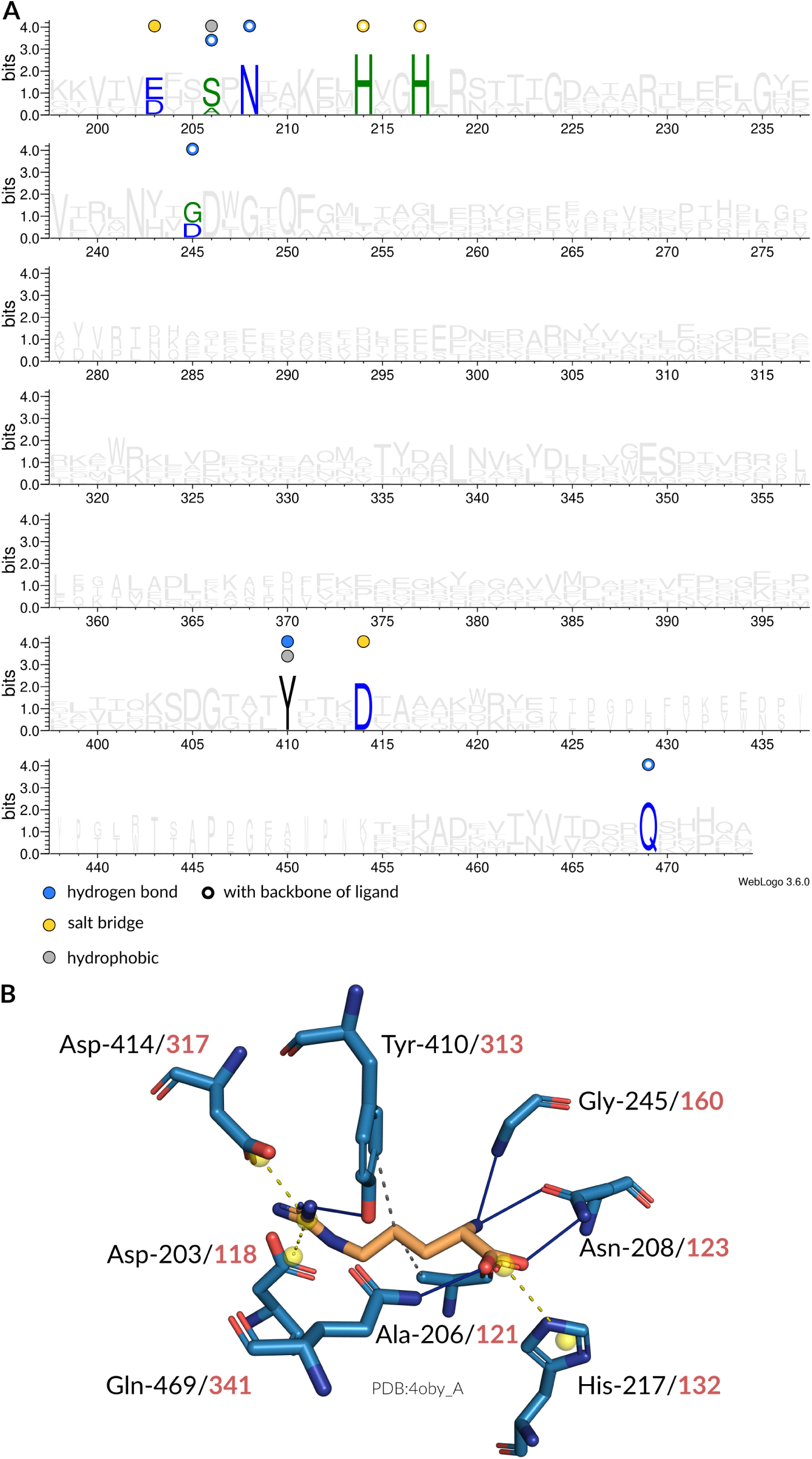
Interaction patterns of arginyl-tRNA synthetase (ArgRS). (**A**) Sequence logo (2) of representative sequences for ArgRSs. Non-covalent interactions with the amino acid ligand occurring at certain positions are indicated by colored circles. Filled circles are interactions with the side chain atoms, while hollow circles are interactions with any of the backbone atoms of the amino acid ligand. (**B**) Depiction of interactions in the binding site (blue stick model) of an ArgRS from *Escherichia coli* (PDB:4oby chain A) with its ligand (orange stick model). Here, hydrogen bonds (solid blue lines), salt bridges (dashed yellow lines) and hydrophobic interactions (dashed gray lines) are established. The sequence positions of the interacting residues are given in accordance to the multiple sequence alignment (MSA) (black) as well as the original structure (red).

**Fig. S5.**
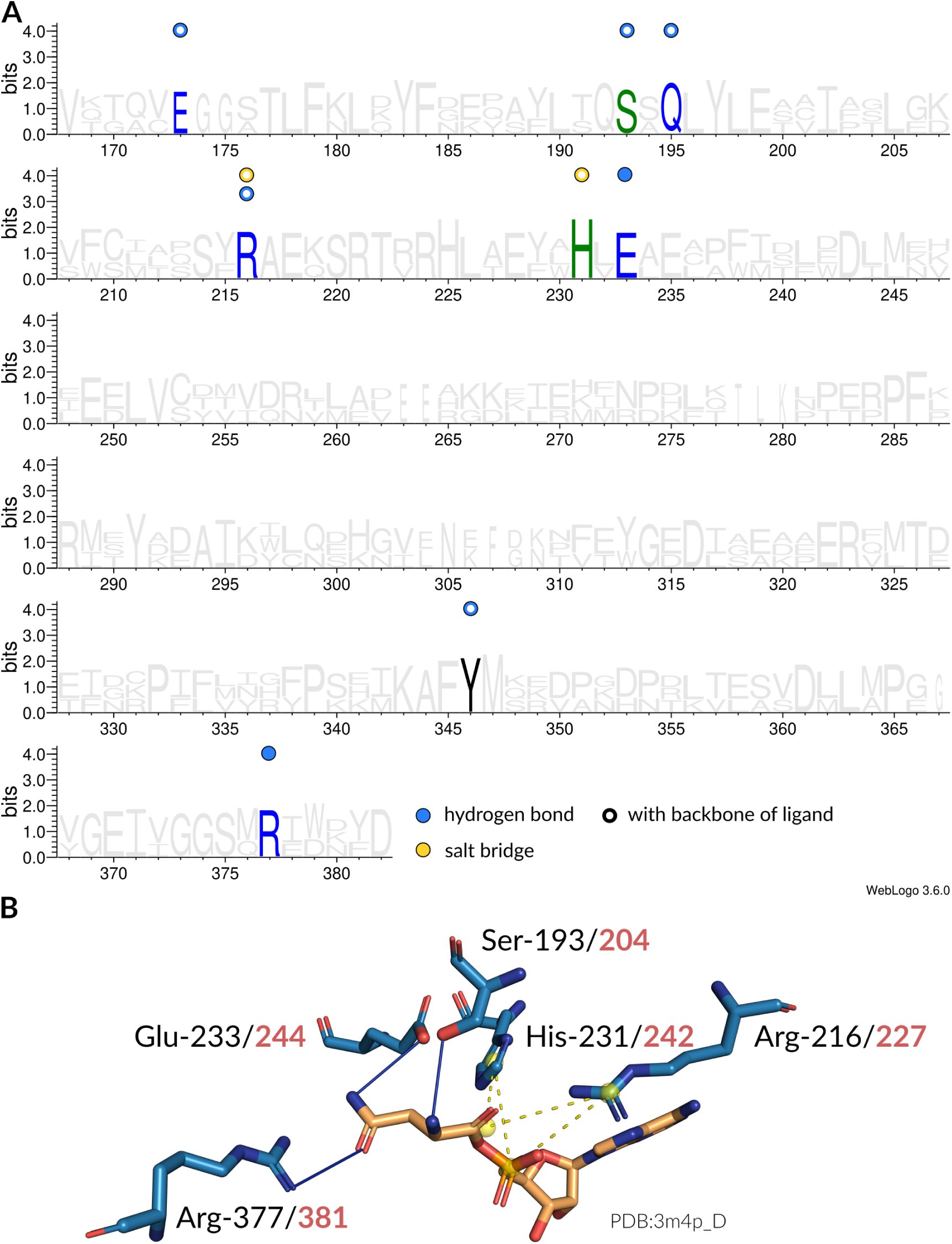
Interaction patterns of asparaginyl-tRNA synthetase (AsnRS). (**A**) Sequence logo (2) of representative sequences for AsnRSs. Non-covalent interactions with the amino acid ligand occurring at certain positions are indicated by colored circles. Filled circles are interactions with the side chain atoms, while hollow circles are interactions with any of the backbone atoms of the amino acid ligand. (**B**) Depiction of interactions in the binding site (blue stick model) of an AsnRS from *Entamoeba histolytica* (PDB:3m4p chain D) with its ligand (orange stick model). Here, hydrogen bonds (solid blue lines), salt bridges (dashed yellow lines), and hydrophobic interactions (dashed gray lines) are established. The sequence positions of the interacting residues are given in accordance to the MSA (black) as well as the original structure (red).

**Fig. S6.**
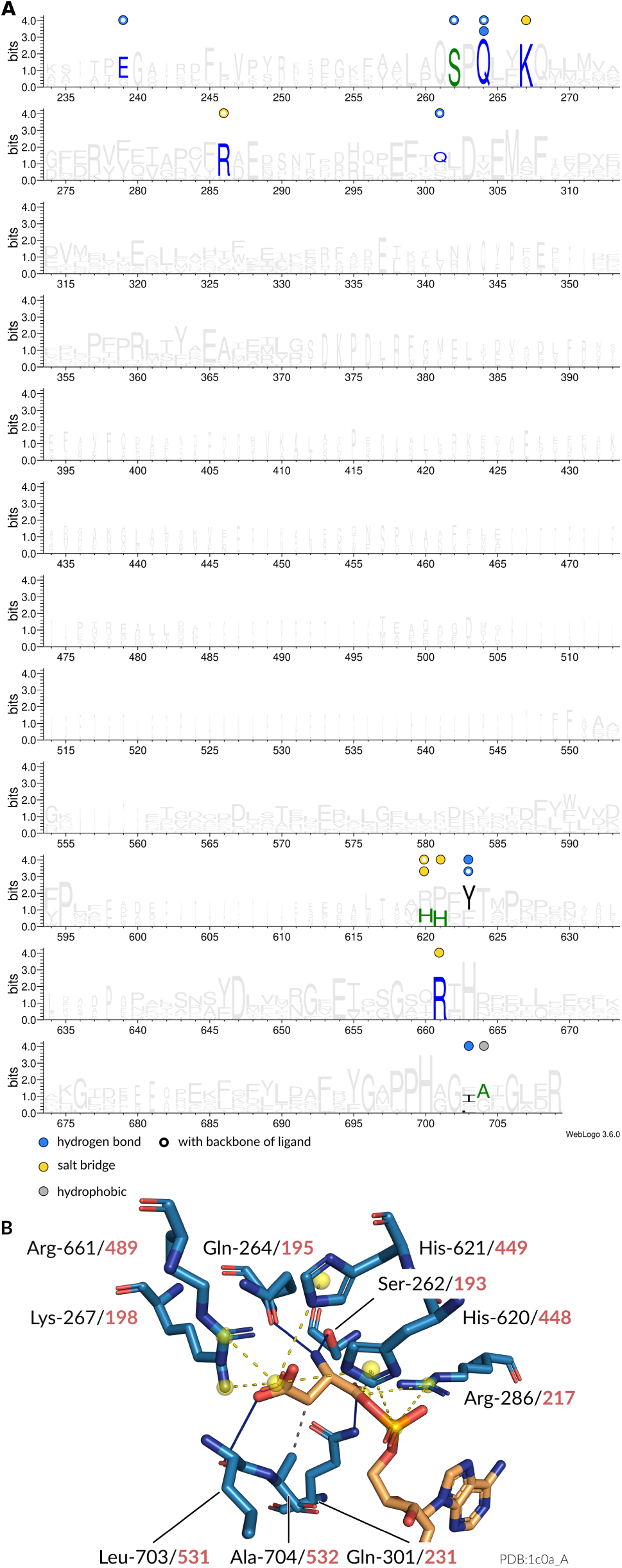
Interaction patterns of aspartyl-tRNA synthetase (AspRS). (**A**) Sequence logo (2) of representative sequences for AspRSs. Non-covalent interactions with the amino acid ligand occurring at certain positions are indicated by colored circles. Filled circles are interactions with the side chain atoms, while hollow circles are interactions with any of the backbone atoms of the amino acid ligand. (**B**) Depiction of interactions in the binding site (blue stick model) of an AspRS from *Escherichia coli* (PDB:1c0a chain A) with its ligand (orange stick model). Here, hydrogen bonds (solid blue lines), salt bridges (dashed yellow lines), and hydrophobic interactions (dashed gray lines) are established. The sequence positions of the interacting residues are given in accordance to the MSA (black) as well as the original structure (red).

**Fig. S7.**
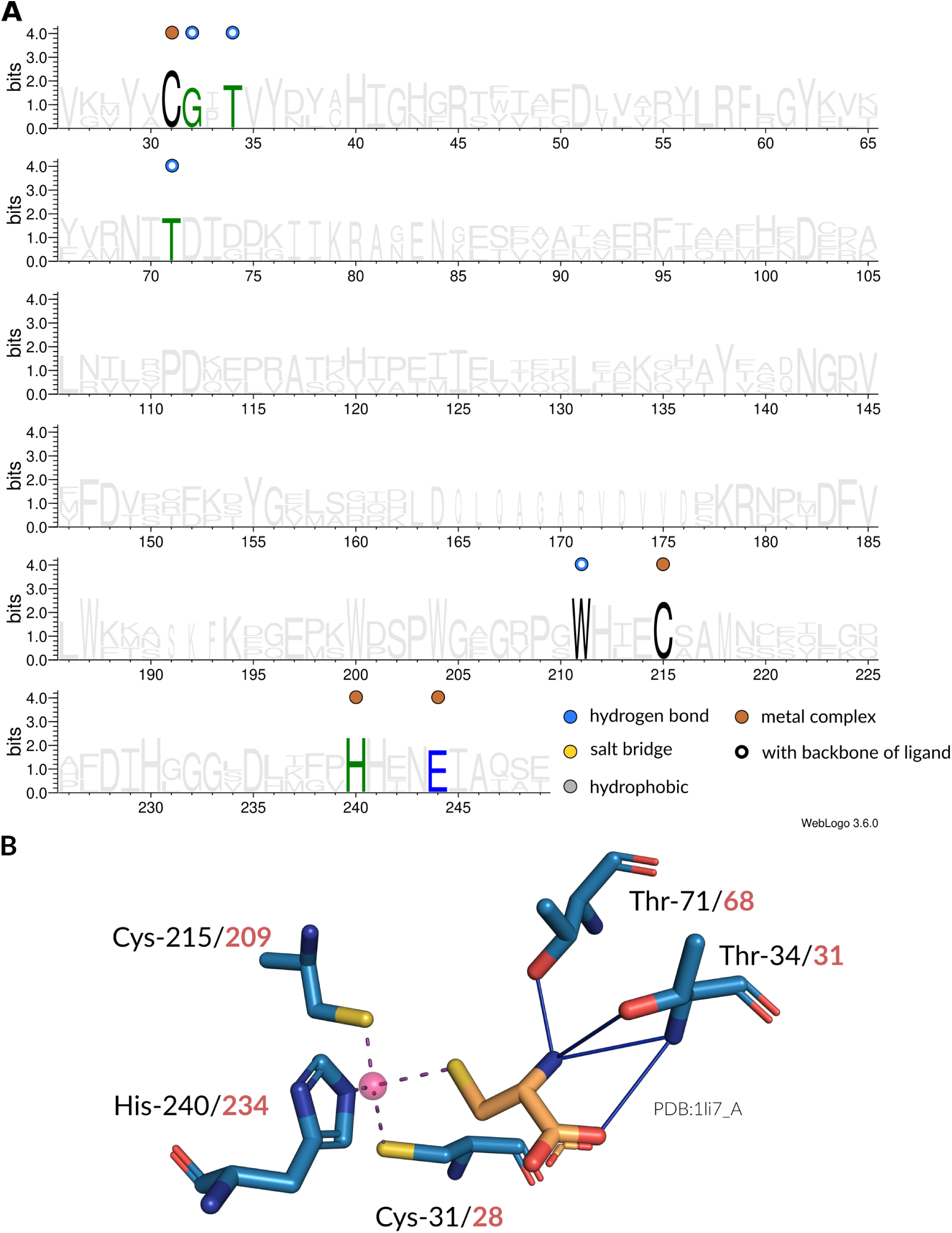
Interaction patterns of cysteinyl-tRNA synthetase (CysRS). (**A**) Sequence logo (2) of representative sequences for CysRSs. Non-covalent interactions with the amino acid ligand occurring at certain positions are indicated by colored circles. Filled circles are interactions with the side chain atoms, while hollow circles are interactions with any of the backbone atoms of the amino acid ligand. (**B**) Depiction of interactions in the binding site (blue stick model) of an CysRS from *Escherichia coli* (PDB:1li7 chain A) with its ligand (orange stick model). Here, hydrogen bonds (solid blue lines) and metal complex interactions (dashed magenta lines) are established. The sequence positions of the interacting residues are given in accordance to the MSA (black) as well as the original structure (red).

**Fig. S8.**
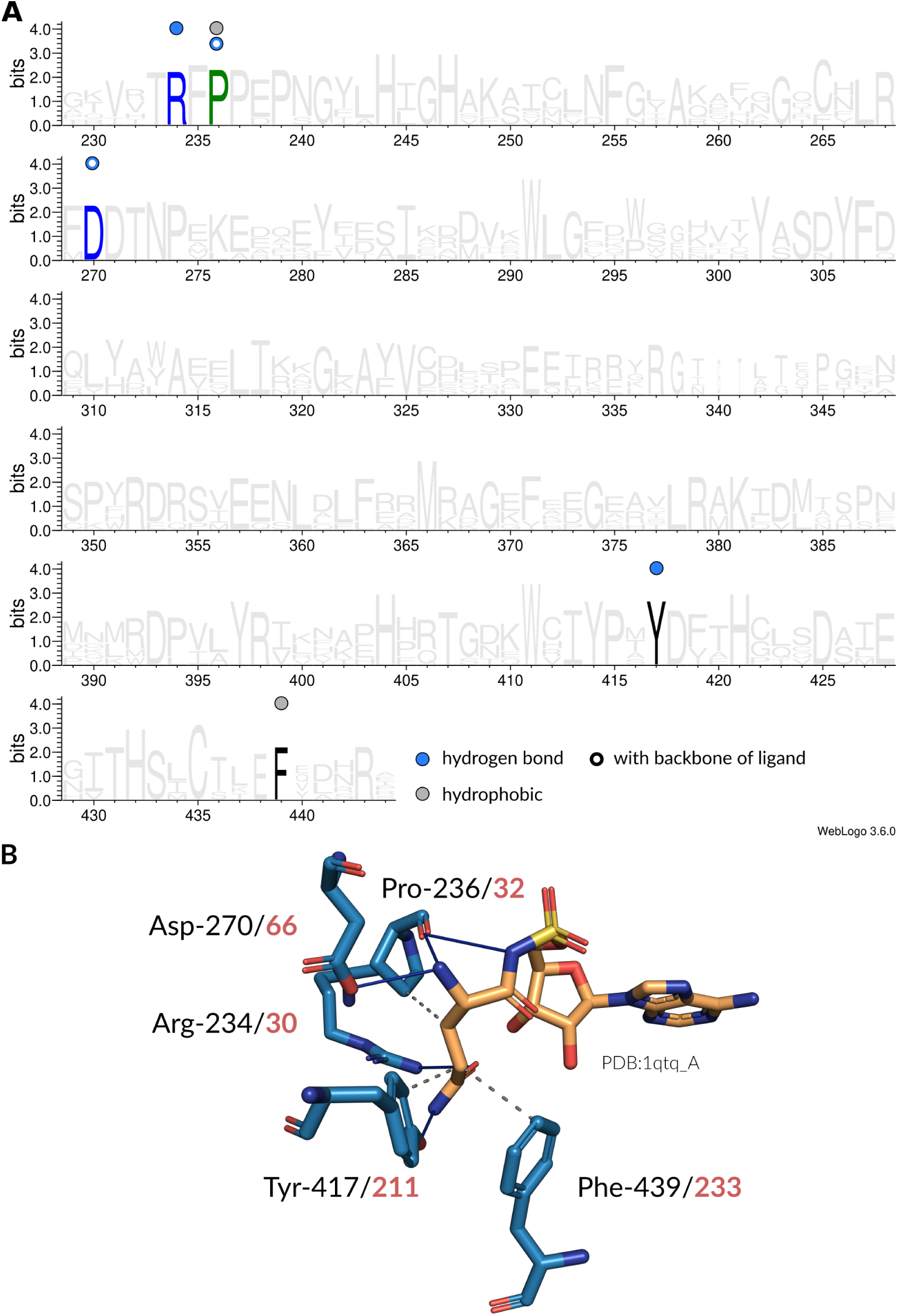
Interaction patterns of glutaminyl-tRNA synthetase (GlnRS). (**A**) Sequence logo (2) of representative sequences for GlnRSs. Non-covalent interactions with the amino acid ligand occurring at certain positions are indicated by colored circles. Filled circles are interactions with the side chain atoms, while hollow circles are interactions with any of the backbone atoms of the amino acid ligand. (**B**) Depiction of interactions in the binding site (blue stick model) of an GlnRS from *Escherichia coli* (PDB:1qtq chain A) with its ligand (orange stick model). Here, hydrogen bonds (solid blue lines) and hydrophobic interactions (dashed gray lines) are established. The sequence positions of the interacting residues are given in accordance to the MSA (black) as well as the original structure (red).

**Fig. S9.**
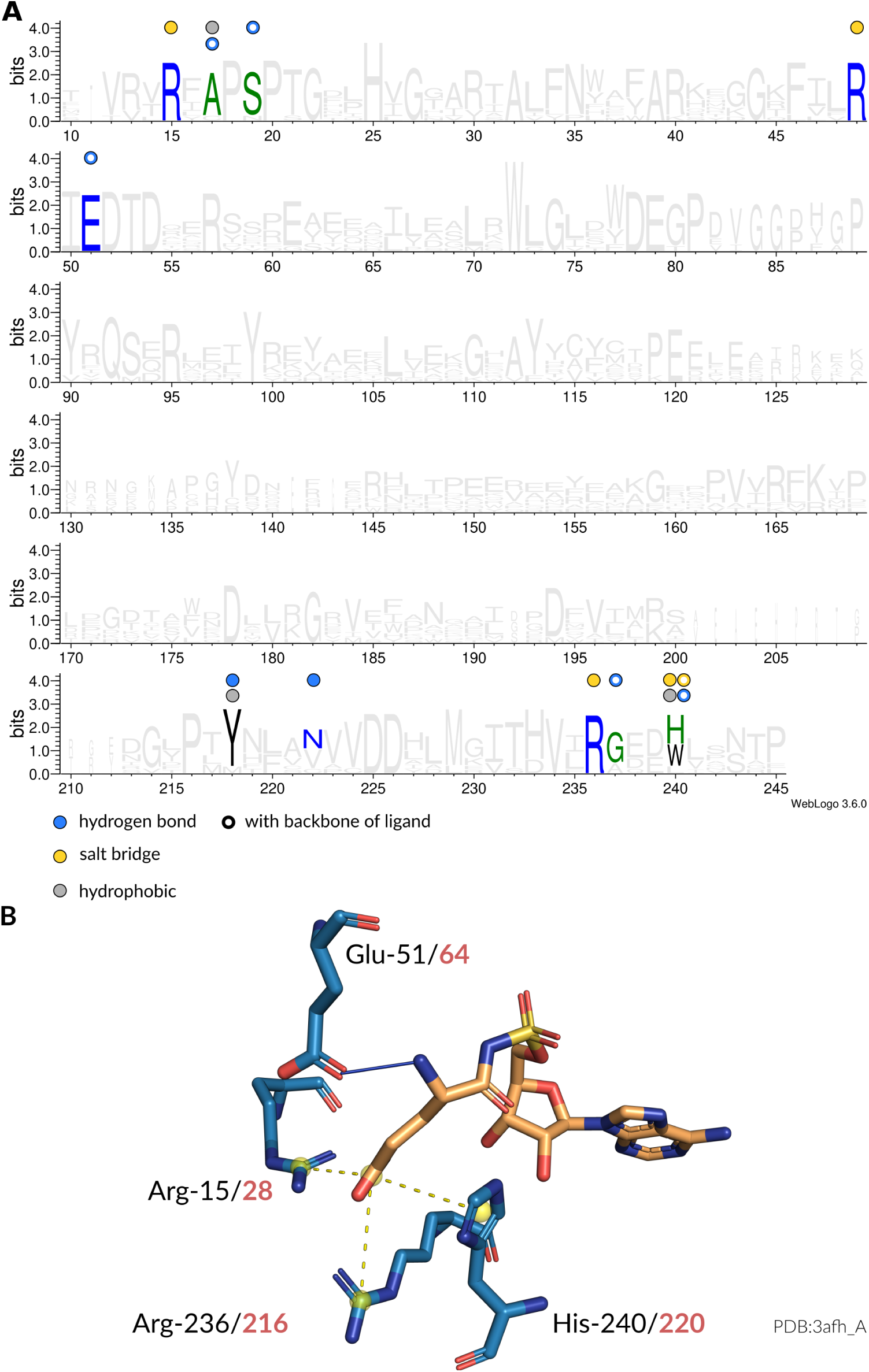
Interaction patterns of glutamyl-tRNA synthetase (GluRS). (**A**) Sequence logo (2) of representative sequences for GluRSs. Non-covalent interactions with the amino acid ligand occurring at certain positions are indicated by colored circles. Filled circles are interactions with the side chain atoms, while hollow circles are interactions with any of the backbone atoms of the amino acid ligand. (**B**) Depiction of interactions in the binding site (blue stick model) of an GluRS from *Thermotoga maritima* (PDB:3afh chain A) with its ligand (orange stick model). Here, hydrogen bonds (solid blue lines) and salt bridges (dashed yellow lines) are established. The sequence positions of the interacting residues are given in accordance to the MSA (black) as well as the original structure (red).

**Fig. S10.**
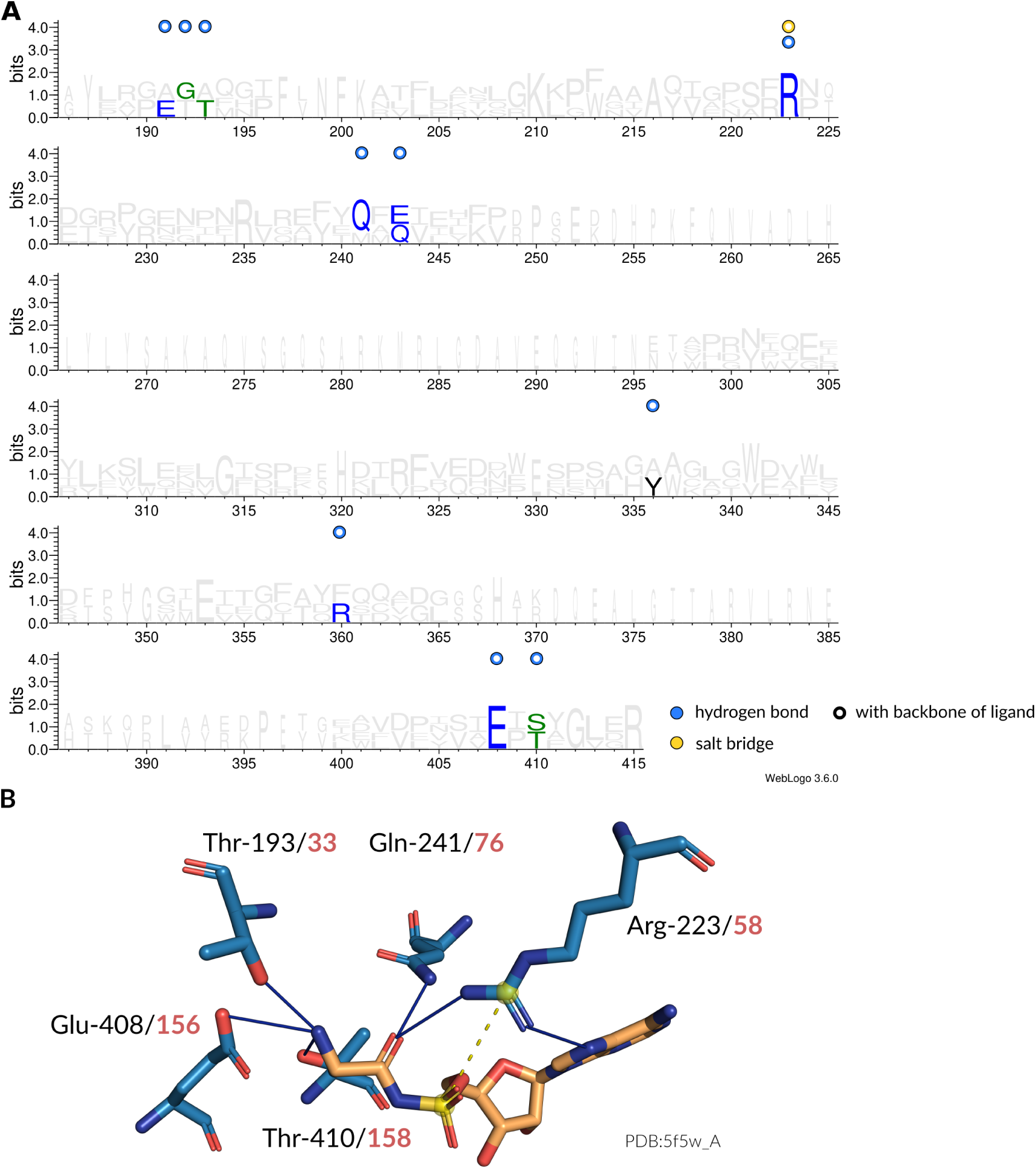
Interaction patterns of glycyl-tRNA synthetase (GlyRS). (**A**) Sequence logo (2) of representative sequences for GlyRSs. Non-covalent interactions with the amino acid ligand occurring at certain positions are indicated by colored circles. Filled circles are interactions with the side chain atoms, while hollow circles are interactions with any of the backbone atoms of the amino acid ligand. (**B**) Depiction of interactions in the binding site (blue stick model) of an GlyRS from *Aquifex aeolicus* (PDB:5f5w chain A) with its ligand (orange stick model). Here, hydrogen bonds (solid blue lines) and salt bridges (dashed yellow lines) are established. The sequence positions of the interacting residues are given in accordance to the MSA (black) as well as the original structure (red).

**Fig. S11.**
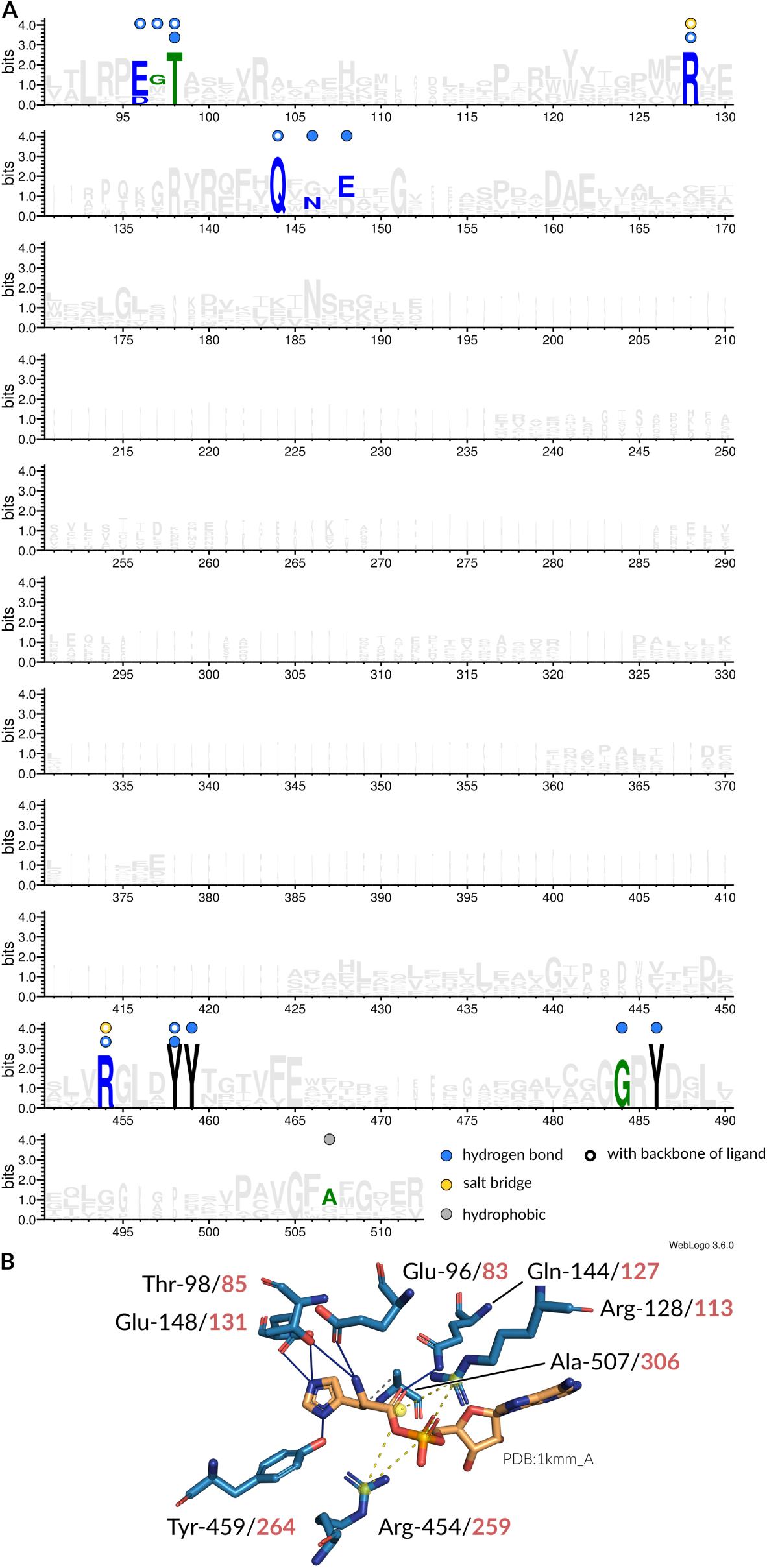
Interaction patterns of histidyl-tRNA synthetase (HisRS). (**A**) Sequence logo (2) of representative sequences for HisRSs. Non-covalent interactions with the amino acid ligand occurring at certain positions are indicated by colored circles. Filled circles are interactions with the side chain atoms, while hollow circles are interactions with any of the backbone atoms of the amino acid ligand. (**B**) Depiction of interactions in the binding site (blue stick model) of an HisRS from *Escherichia coli* (PDB:1kmm chain A) with its ligand (orange stick model). Here, hydrogen bonds (solid blue lines), salt bridges (dashed yellow lines), and hydrophobic interactions (dashed gray lines) are established. The sequence positions of the interacting residues are given in accordance to the MSA (black) as well as the original structure (red).

**Fig. S12.**
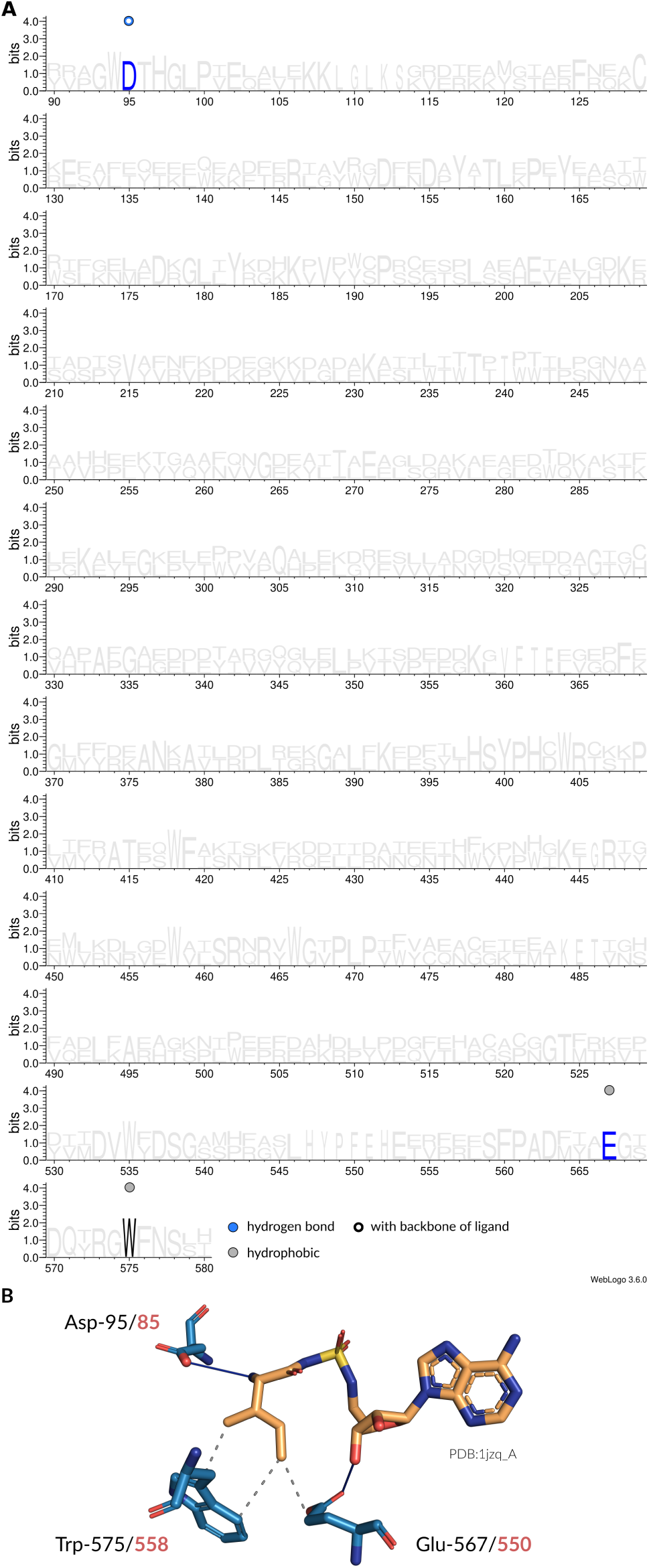
Interaction patterns of isoleucyl-tRNA synthetase (IleRS). (**A**) Sequence logo (2) of representative sequences for IleRSs. Non-covalent interactions with the amino acid ligand occurring at certain positions are indicated by colored circles. Filled circles are interactions with the side chain atoms, while hollow circles are interactions with any of the backbone atoms of the amino acid ligand. (**B**) Depiction of interactions in the binding site (blue stick model) of an IleRS from *Thermus thermophilus* (PDB:1jzq chain A) with its ligand (orange stick model). Here, hydrogen bonds (solid blue lines) and hydrophobic interactions (dashed gray lines) are established. The sequence positions of the interacting residues are given in accordance to the MSA (black) as well as the original structure (red).

**Fig. S13.**
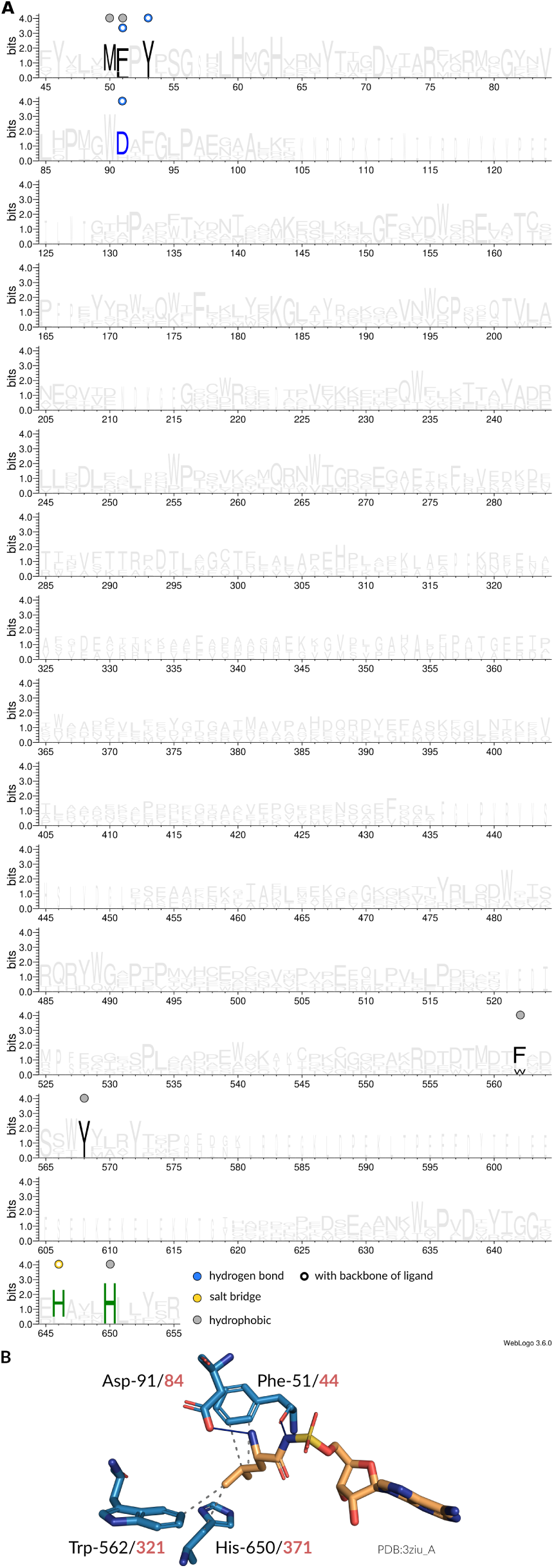
Interaction patterns of leucyl-tRNA synthetase (LeuRS). (**A**) Sequence logo (2) of representative sequences for LeuRSs. Non-covalent interactions with the amino acid ligand occurring at certain positions are indicated by colored circles. Filled circles are interactions with the side chain atoms, while hollow circles are interactions with any of the backbone atoms of the amino acid ligand. (**B**) Depiction of interactions in the binding site (blue stick model) of an LeuRS from *Mycoplasma mobile* (PDB:3ziu chain A) with its ligand (orange stick model). Here, hydrogen bonds (solid blue lines) and hydrophobic interactions (dashed gray lines) are established. The sequence positions of the interacting residues are given in accordance to the MSA (black) as well as the original structure (red).

**Fig. S14.**
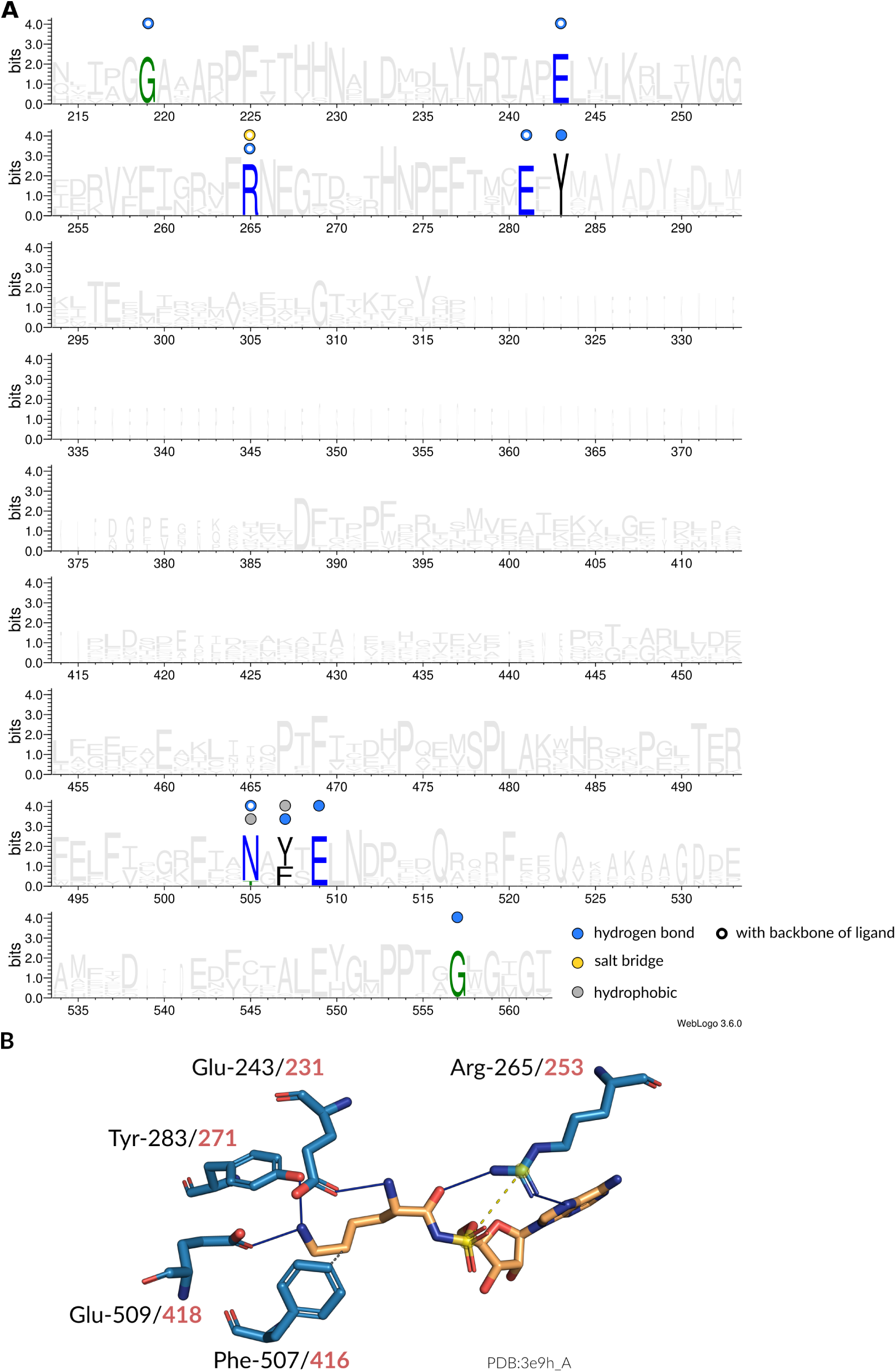
Interaction patterns of lysyl-tRNA synthetase (LysRS). (**A**) Sequence logo (2) of representative sequences for LysRSs. Non-covalent interactions with the amino acid ligand occurring at certain positions are indicated by colored circles. Filled circles are interactions with the side chain atoms, while hollow circles are interactions with any of the backbone atoms of the amino acid ligand. (**B**) Depiction of interactions in the binding site (blue stick model) of an LysRS from *Geobacillus stearothermophilus* (PDB:3e9h chain A) with its ligand (orange stick model). Here, hydrogen bonds (solid blue lines), salt bridges (dashed yellow lines), and hydrophobic interactions (dashed gray lines) are established. The sequence positions of the interacting residues are given in accordance to the MSA (black) as well as the original structure (red).

**Fig. S15.**
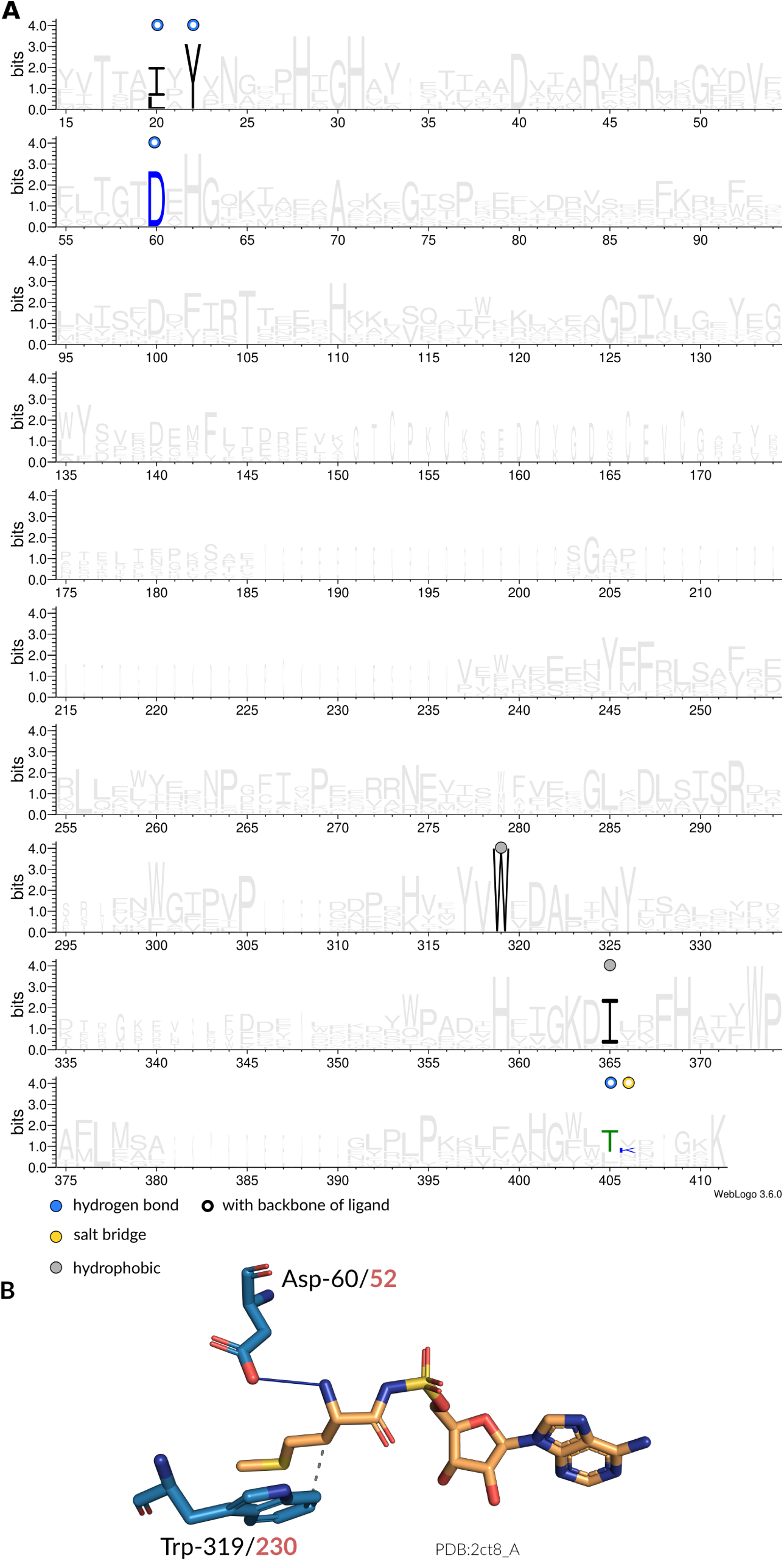
Interaction patterns of methionyl-tRNA synthetase (MetRS). (**A**) Sequence logo (2) of representative sequences for MetRSs. Non-covalent interactions with the amino acid ligand occurring at certain positions are indicated by colored circles. Filled circles are interactions with the side chain atoms, while hollow circles are interactions with any of the backbone atoms of the amino acid ligand. (**B**) Depiction of interactions in the binding site (blue stick model) of an MetRS from *Aquifex aeolicus* (PDB:2ct8 chain A) with its ligand (orange stick model). Here, hydrogen bonds (solid blue lines) and hydrophobic interactions (dashed gray lines) are established. The sequence positions of the interacting residues are given in accordance to the MSA (black) as well as the original structure (red).

**Fig. S16.**
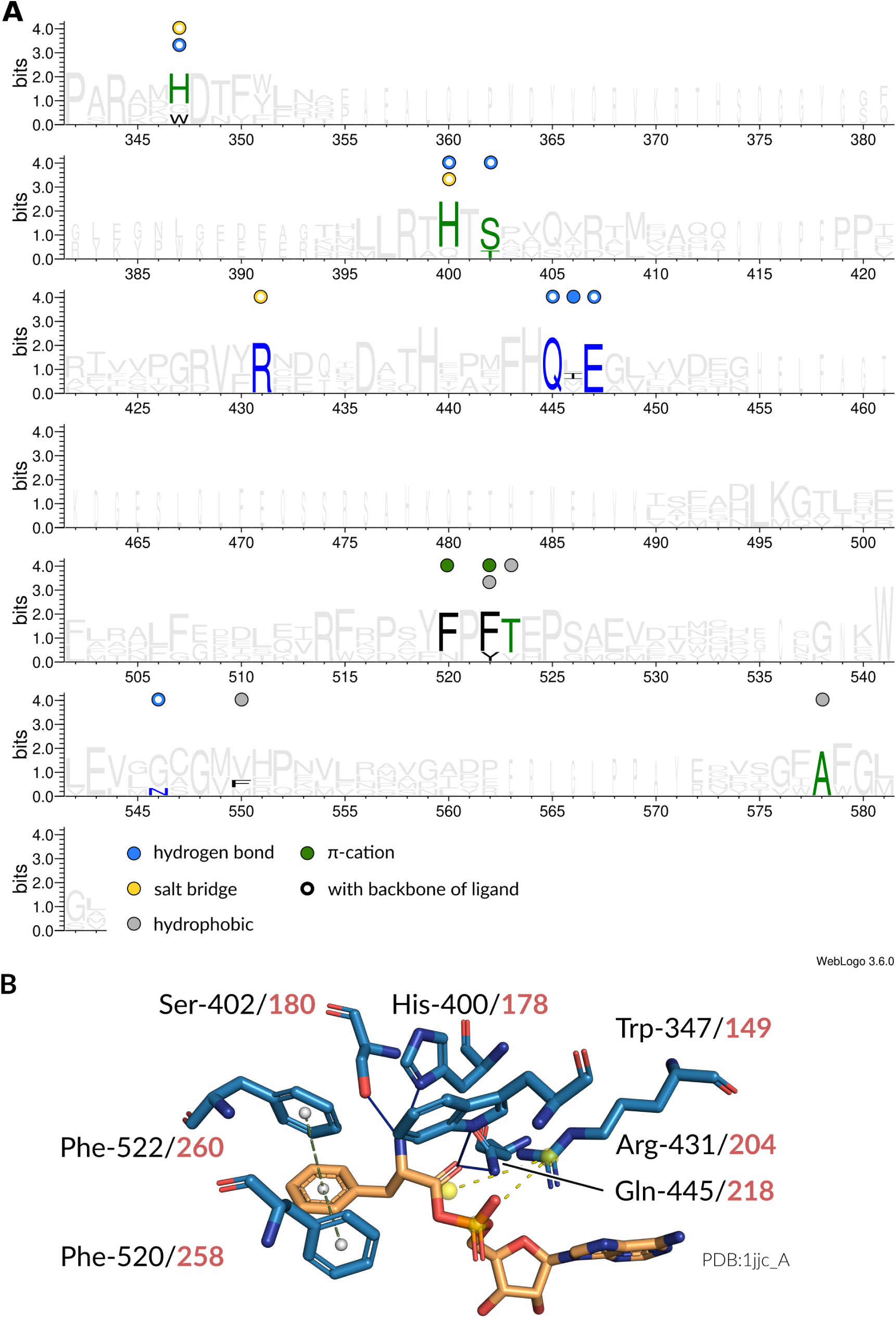
Interaction patterns of phenylalanine-tRNA synthetase (PheRS). (**A**) Sequence logo (2) of representative sequences for PheRSs. Non-covalent interactions with the amino acid ligand occurring at certain positions are indicated by colored circles. Filled circles are interactions with the side chain atoms, while hollow circles are interactions with any of the backbone atoms of the amino acid ligand. (**B**) Depiction of interactions in the binding site (blue stick model) of an PheRS from *Thermus thermophilus* (PDB:1jjc chain A) with its ligand (orange stick model). Here, hydrogen bonds (solid blue lines), salt bridges (dashed yellow lines), and *π*-stacking interactions (dashed green lines) are established. The sequence positions of the interacting residues are given in accordance to the MSA (black) as well as the original structure (red).

**Fig. S17.**
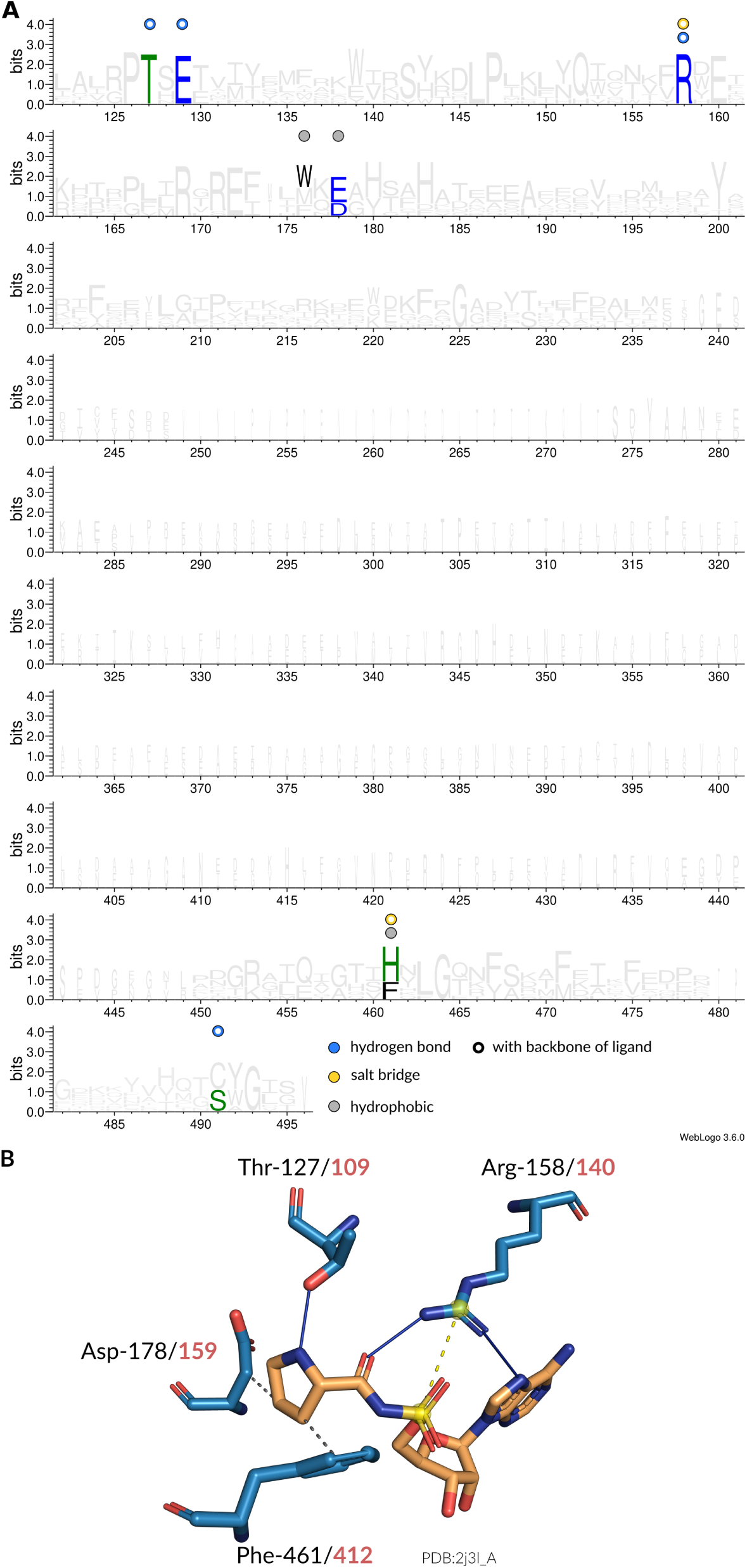
Interaction patterns of prolyl-tRNA synthetase (ProRS). (**A**) Sequence logo (2) of representative sequences for ProRSs. Non-covalent interactions with the amino acid ligand occurring at certain positions are indicated by colored circles. Filled circles are interactions with the side chain atoms, while hollow circles are interactions with any of the backbone atoms of the amino acid ligand. (**B**) Depiction of interactions in the binding site (blue stick model) of an ProRS from *Enterococcus faecalis* (PDB:2j3l chain A) with its ligand (orange stick model). Here, hydrogen bonds (solid blue lines), salt bridges (dashed yellow lines), and hydrophobic interactions (dashed gray lines) are established. The sequence positions of the interacting residues are given in accordance to the MSA (black) as well as the original structure (red).

**Fig. S18.**
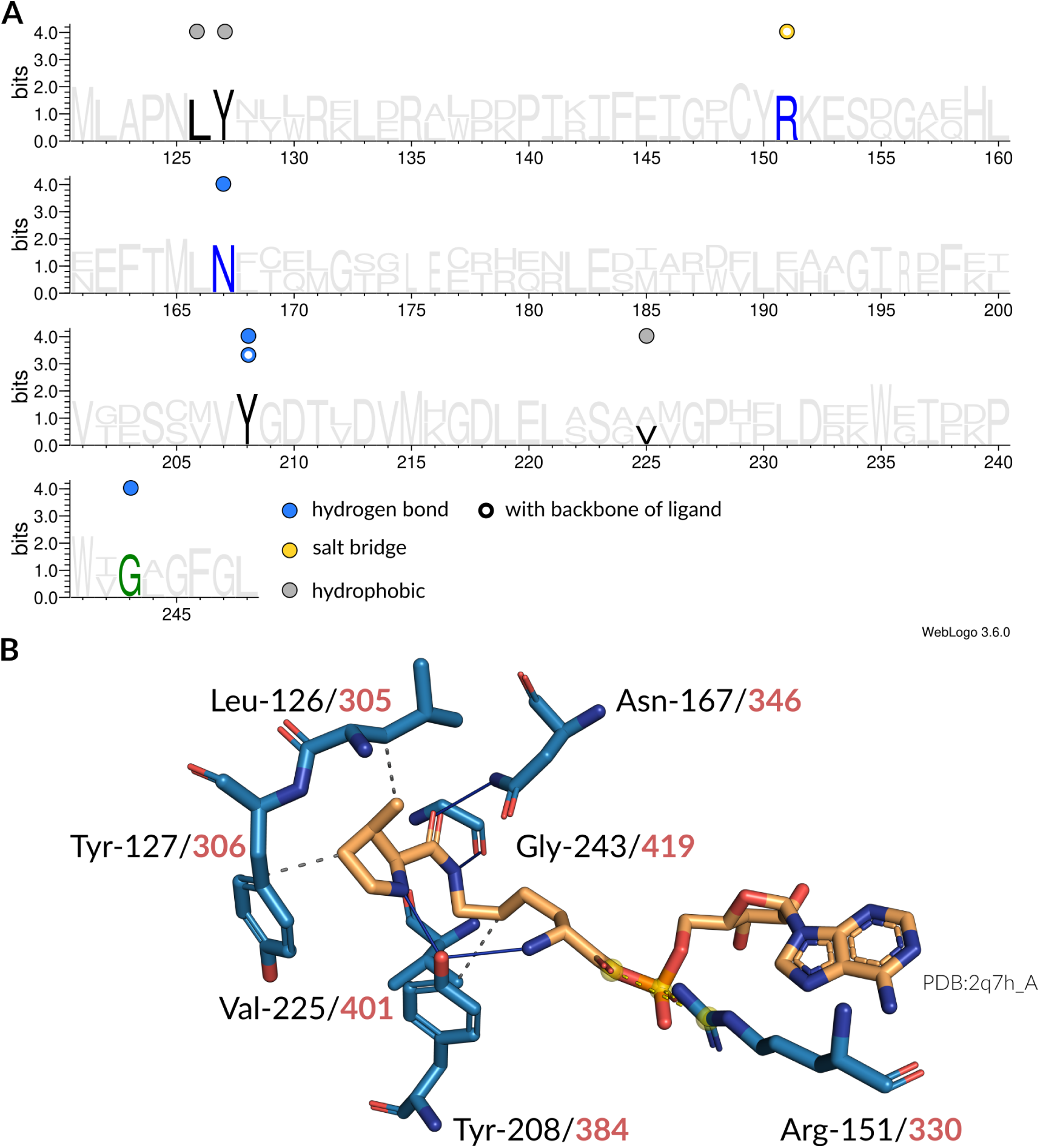
Interaction patterns of pyrrolysyl-tRNA synthetase (PylRS). (**A**) Sequence logo (2) of representative sequences for PylRSs. Non-covalent interactions with the amino acid ligand occurring at certain positions are indicated by colored circles. Filled circles are interactions with the side chain atoms, while hollow circles are interactions with any of the backbone atoms of the amino acid ligand. (**B**) Depiction of interactions in the binding site (blue stick model) of an PylRS from *Methanosarcina mazei* (PDB:2q7h chain A) with its ligand (orange stick model). Here, hydrogen bonds (solid blue lines), salt bridges (dashed yellow lines), and hydrophobic interactions (dashed gray lines) are established. The sequence positions of the interacting residues are given in accordance to the MSA (black) as well as the original structure (red).

**Fig. S19.**
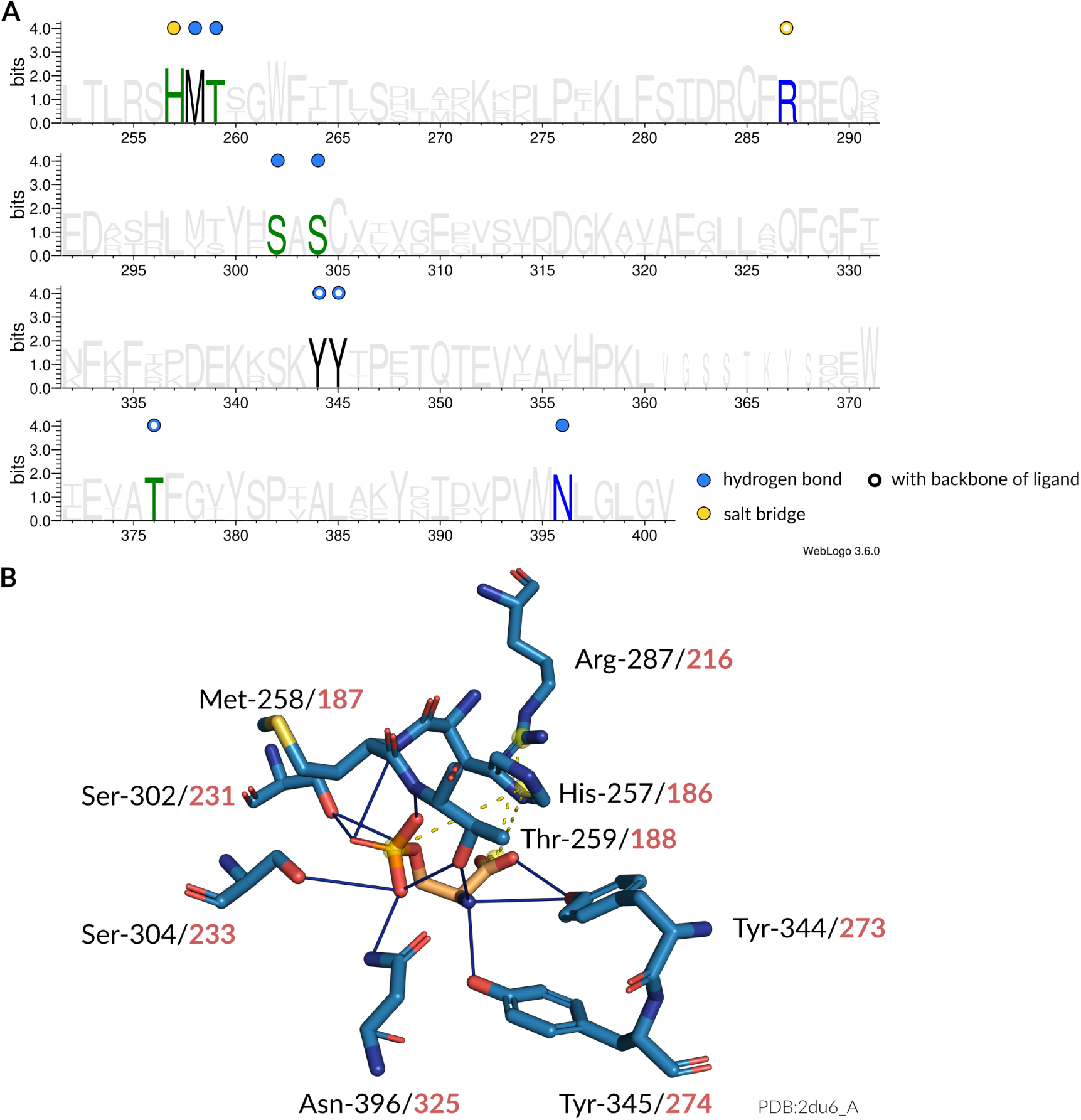
Interaction patterns of phosphoseryl-tRNA synthetase (SepRS). (**A**) Sequence logo (2) of representative sequences for SepRSs. Non-covalent interactions with the amino acid ligand occurring at certain positions are indicated by colored circles. Filled circles are interactions with the side chain atoms, while hollow circles are interactions with any of the backbone atoms of the amino acid ligand. (**B**) Depiction of interactions in the binding site (blue stick model) of an SepRS from *Archaeoglobus fulgidus* (PDB:2du6 chain A) with its ligand (orange stick model). Here, hydrogen bonds (solid blue lines) and salt bridges (dashed yellow lines) are established. The sequence positions of the interacting residues are given in accordance to the MSA (black) as well as the original structure (red).

**Fig. S20.**
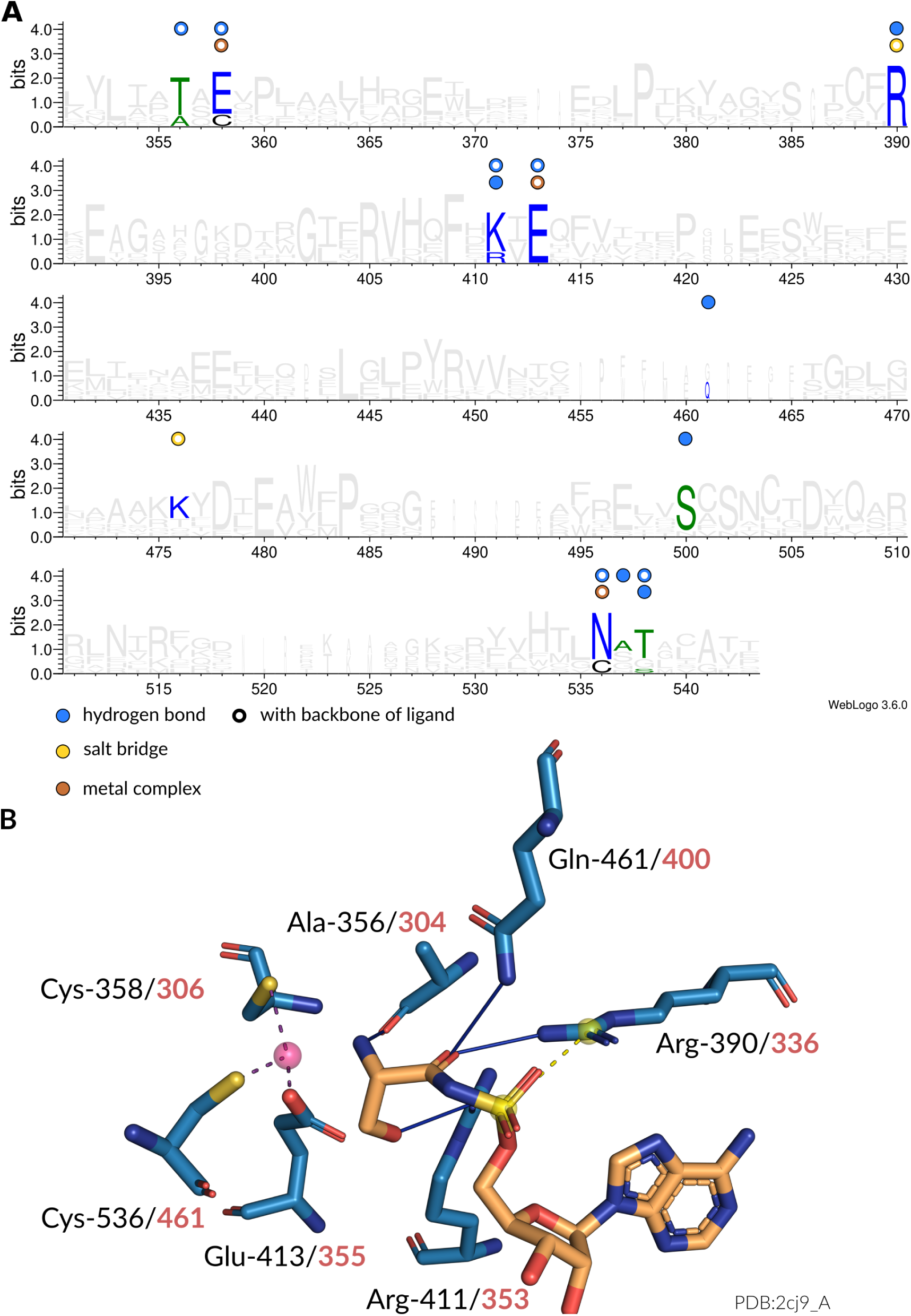
Interaction patterns of seryl-tRNA synthetase (SerRS). (**A**) Sequence logo (2) of representative sequences for SerRSs. Non-covalent interactions with the amino acid ligand occurring at certain positions are indicated by colored circles. Filled circles are interactions with the side chain atoms, while hollow circles are interactions with any of the backbone atoms of the amino acid ligand. (**B**) Depiction of interactions in the binding site (blue stick model) of an SerRS from *Methanosarcina barkeri* (PDB:2cj9 chain A) with its ligand (orange stick model). Here, hydrogen bonds (solid blue lines), salt bridges (dashed yellow lines), and metal complex interactions (dashed magenta lines) are established. The sequence positions of the interacting residues are given in accordance to the MSA (black) as well as the original structure (red).

**Fig. S21.**
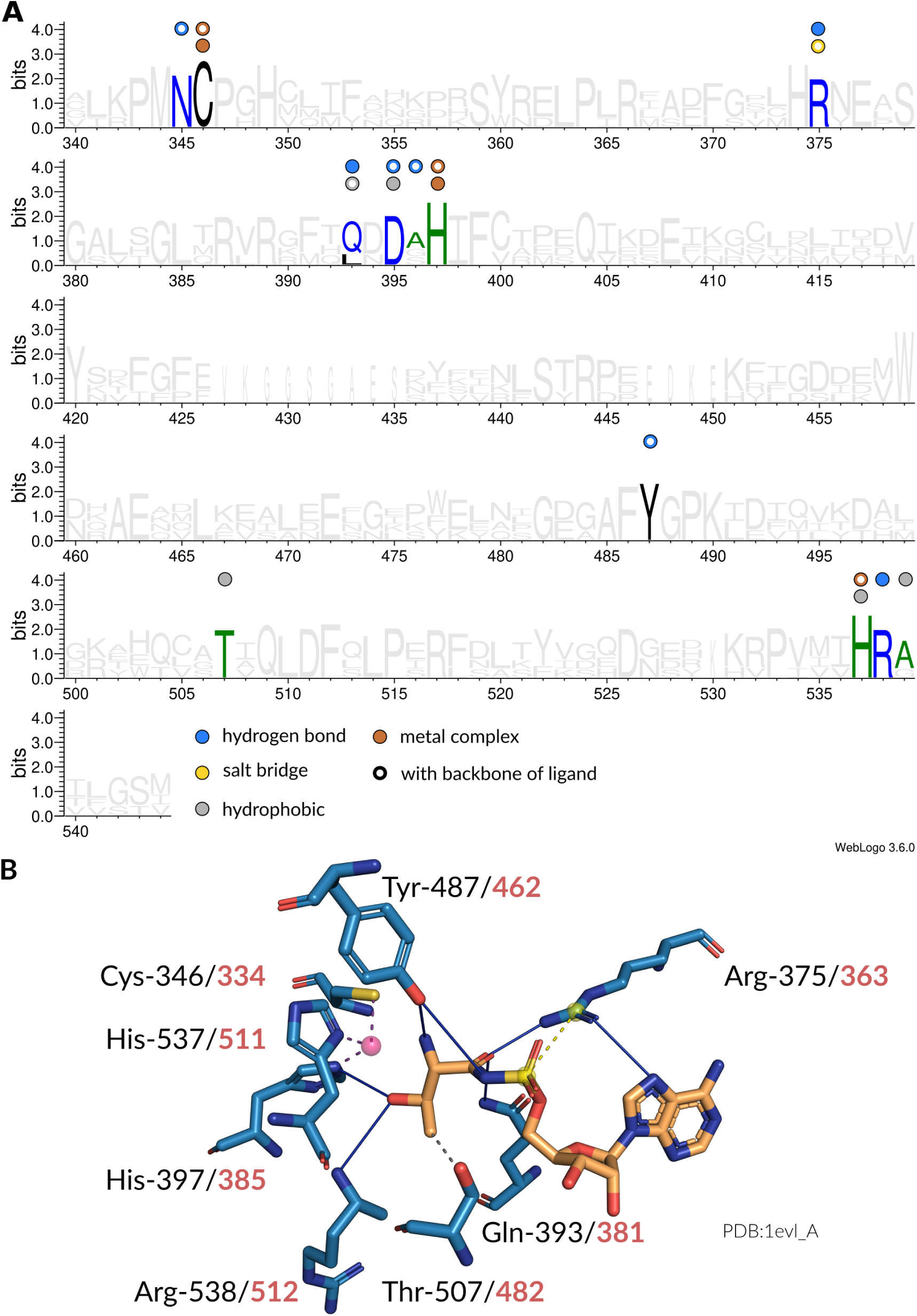
Interaction patterns of threonyl-tRNA synthetase (ThrRS). (**A**) Sequence logo (2) of representative sequences for ThrRSs. Non-covalent interactions with the amino acid ligand occurring at certain positions are indicated by colored circles. Filled circles are interactions with the side chain atoms, while hollow circles are interactions with any of the backbone atoms of the amino acid ligand. (**B**) Depiction of interactions in the binding site (blue stick model) of an ThrRS from *Escherichia coli* (PDB:1evl chain A) with its ligand (orange stick model). Here, hydrogen bonds (solid blue lines), salt bridges (dashed yellow lines), metal complex interactions (dashed magenta lines), and hydrophobic interactions (dashed gray lines) are established. The sequence positions of the interacting residues are given in accordance to the MSA (black) as well as the original structure (red).

**Fig. S22.**
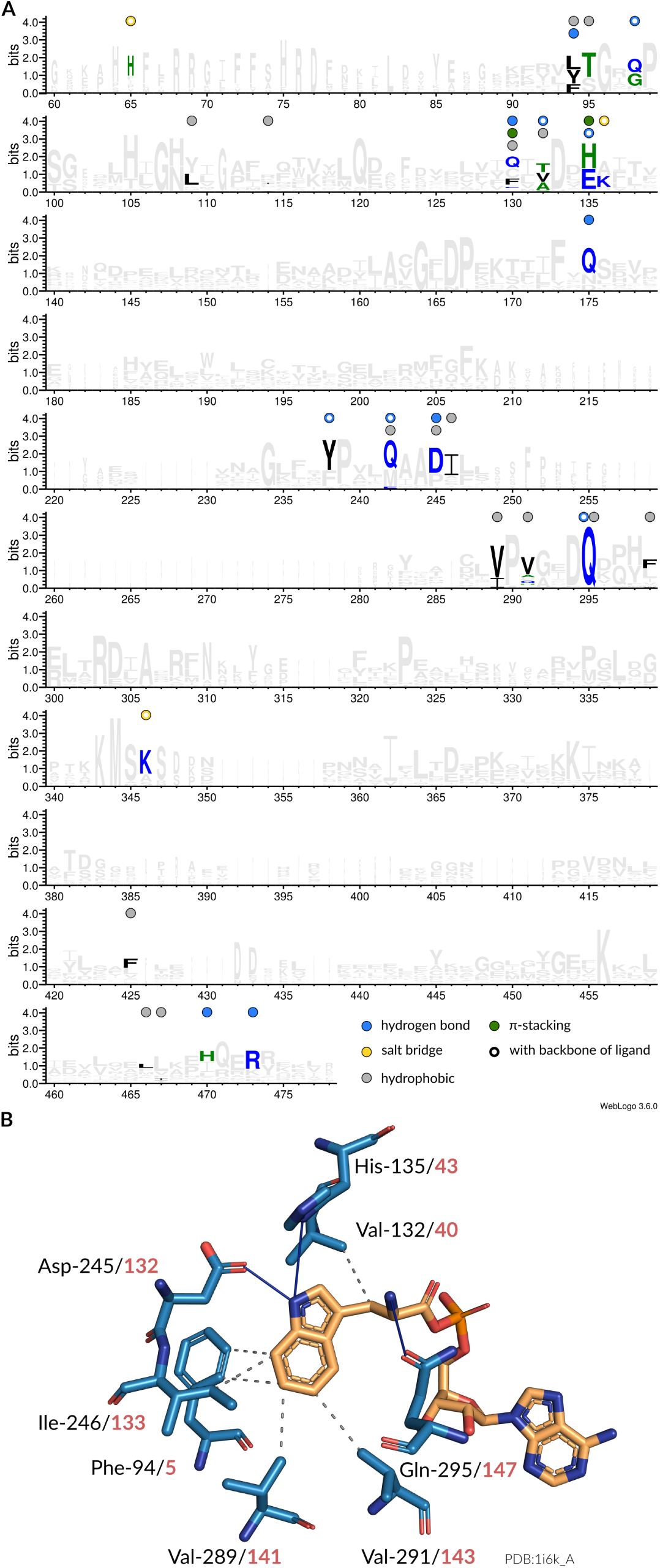
Interaction patterns of tryptophanyl-tRNA synthetase (TrpRS). (**A**) Sequence logo (2) of representative sequences for TrpRSs. Non-covalent interactions with the amino acid ligand occurring at certain positions are indicated by colored circles. Filled circles are interactions with the side chain atoms, while hollow circles are interactions with any of the backbone atoms of the amino acid ligand. (**B**) Depiction of interactions in the binding site (blue stick model) of an TrpRS from *Geobacillus stearothermophilus* (PDB:1i6k chain A) with its ligand (orange stick model). Here, hydrogen bonds (solid blue lines) and hydrophobic interactions (dashed gray lines) are established. The sequence positions of the interacting residues are given in accordance to the MSA (black) as well as the original structure (red).

**Fig. S23.**
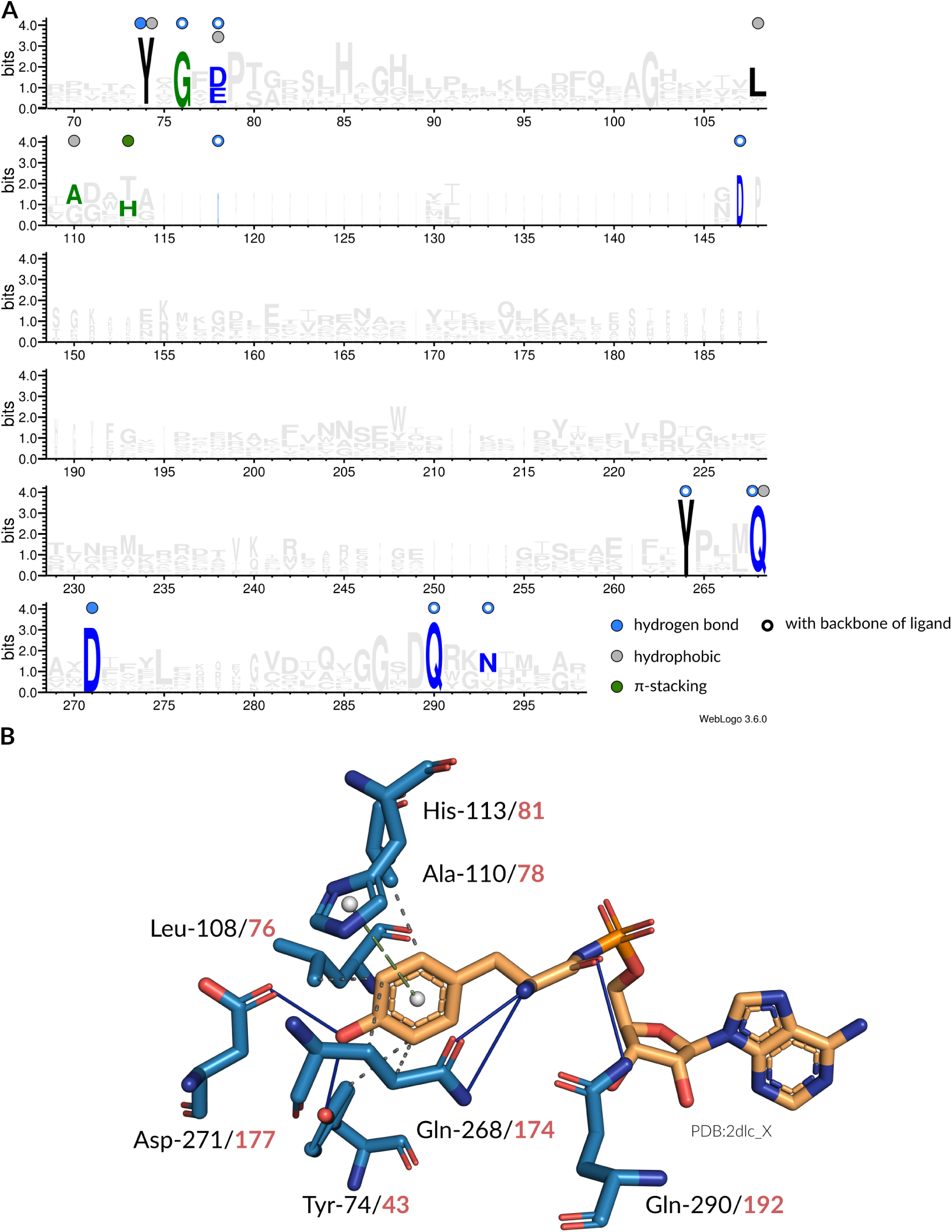
Interaction patterns of tyrosyl-tRNA synthetase (TyrRS). (**A**) Sequence logo (2) of representative sequences for TyrRSs. Non-covalent interactions with the amino acid ligand occurring at certain positions are indicated by colored circles. Filled circles are interactions with the side chain atoms, while hollow circles are interactions with any of the backbone atoms of the amino acid ligand. (**B**) Depiction of interactions in the binding site (blue stick model) of an TyrRS from *Saccharomyces cerevisiae* (PDB:2dlc chain X) with its ligand (orange stick model). Here, hydrogen bonds (solid blue lines), *π*-stacking interactions (dashed greeen lines), and hydrophobic interactions (dashed gray lines) are established. The sequence positions of the interacting residues are given in accordance to the MSA (black) as well as the original structure (red).

**Fig. S24.**
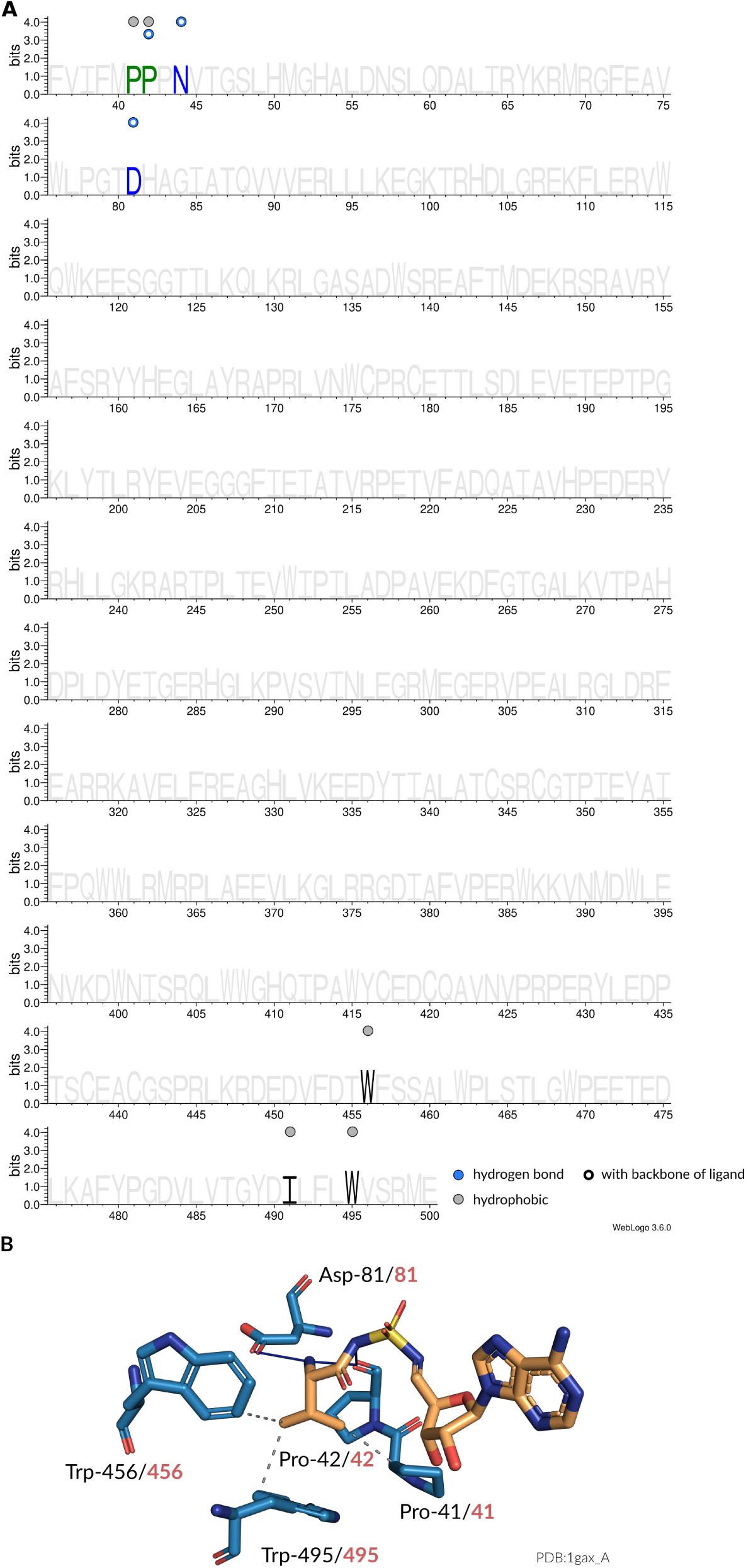
Interaction patterns of valyl-tRNA synthetase (ValRS). (**A**) Sequence logo (2) of representative sequences for ValRSs. Non-covalent interactions with the amino acid ligand occurring at certain positions are indicated by colored circles. Filled circles are interactions with the side chain atoms, while hollow circles are interactions with any of the backbone atoms of the amino acid ligand. (**B**) Depiction of interactions in the binding site (blue stick model) of an ValRS from *Thermus thermophilus* (PDB:1gax chain X) with its ligand (orange stick model). Here, hydrogen bonds (solid blue lines) and hydrophobic interactions (dashed gray lines) are established. The sequence positions of the interacting residues are given in accordance to the MSA (black) as well as the original structure (red).

**Fig. S25.**
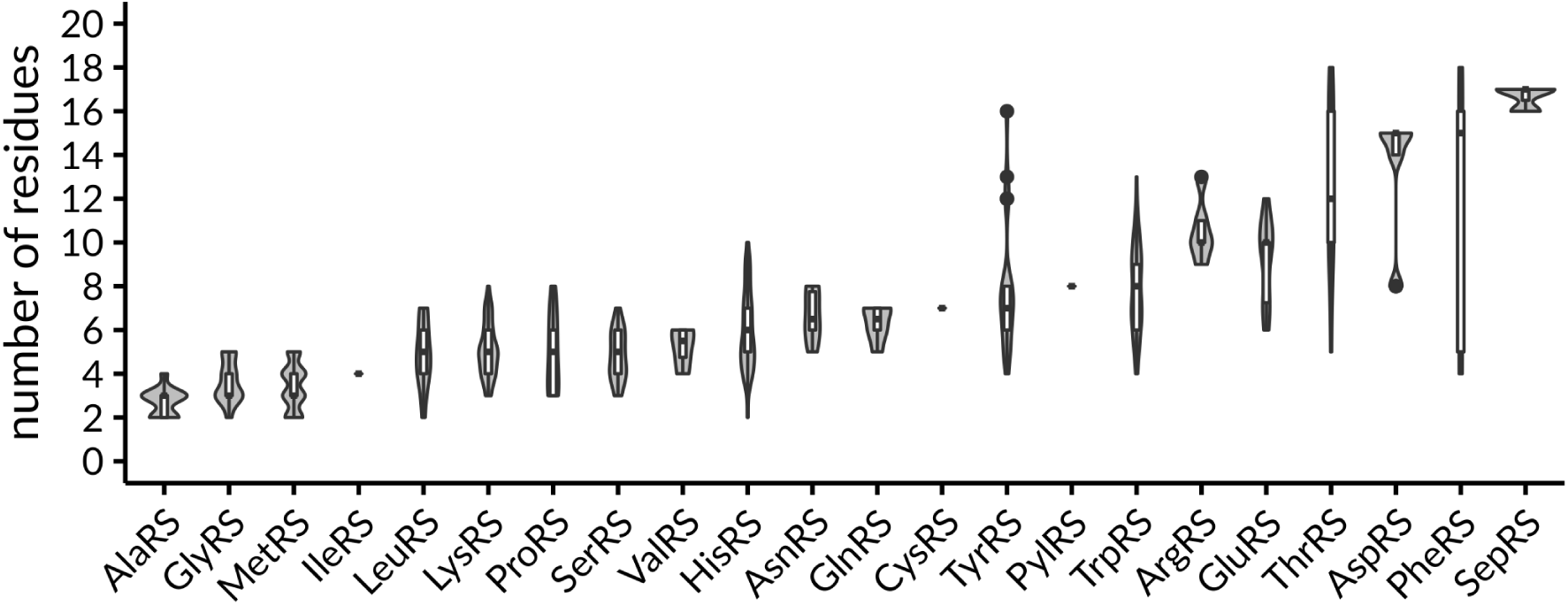
The number of binding site residues involved in specificity-conferring interactions for each aaRSs. Data is sorted by ascending median from left to right.

**Fig. S26.**
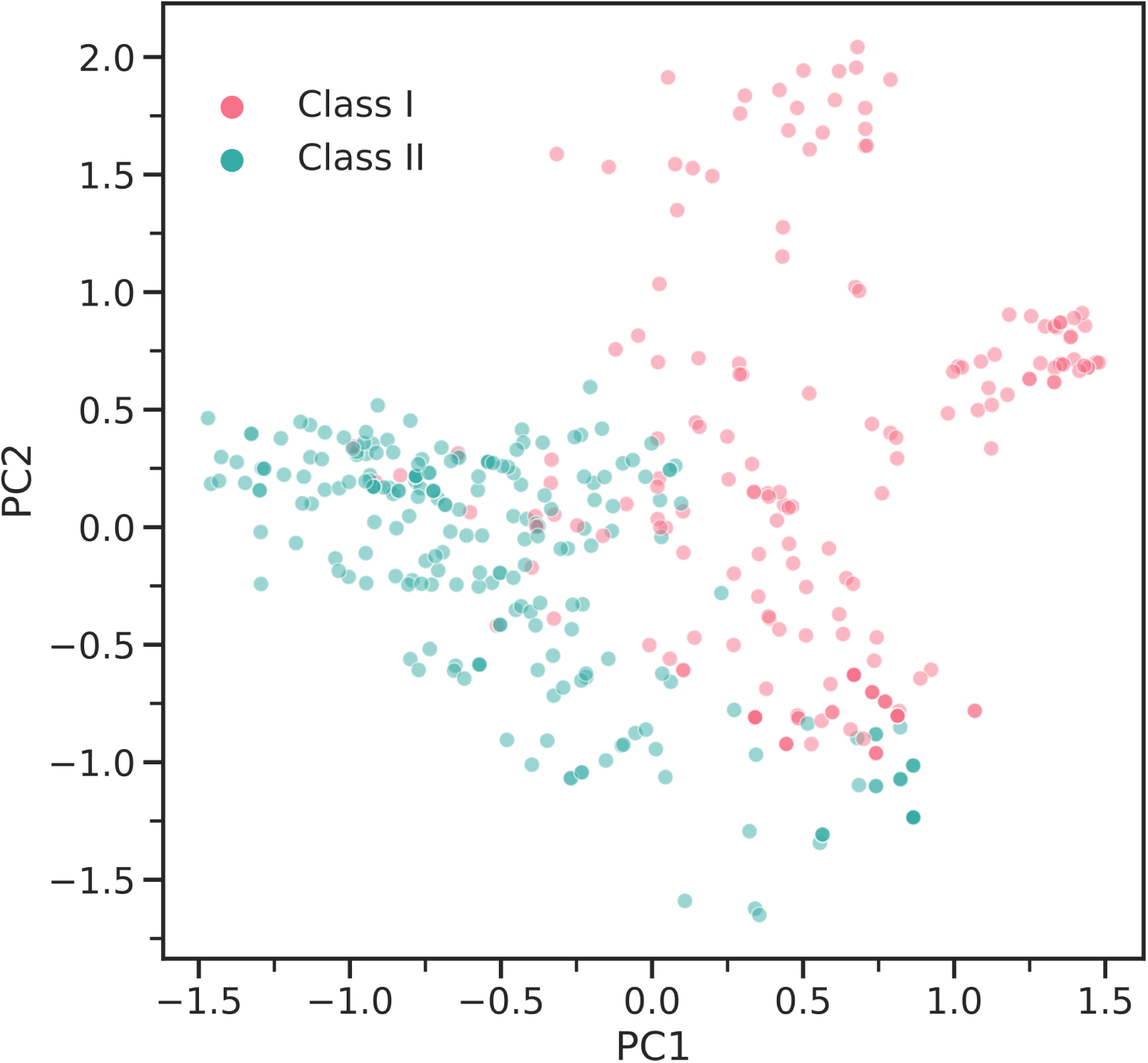
Principal component analysis of interaction fingerprints colored according to the respective aaRS class. The first two components account for 9.24% and 8.44% of the covered variance, respectively. This indicated that the fingerprint representation is high-dimensional abstraction of the complex ligand recognition mechanisms in aaRSs.

**Fig. S27.**
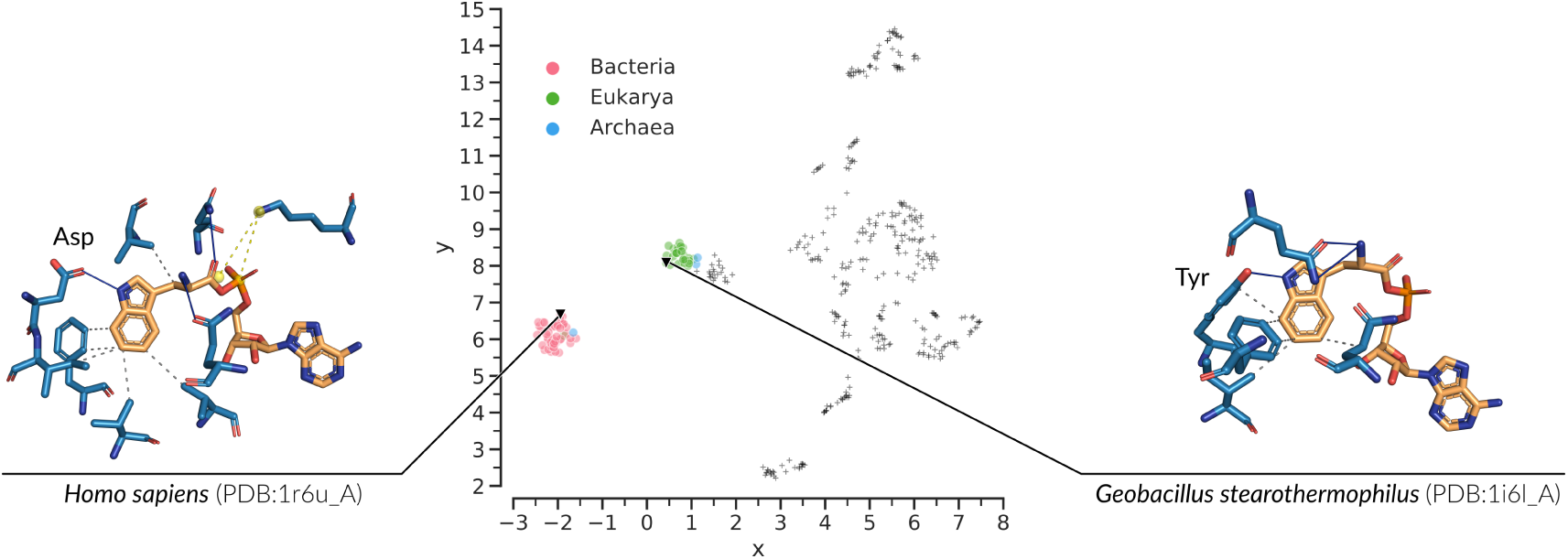
Embedding space of interaction fingerprints. TrpRSs are highlighted and colored by the superkingdom of their species of origin. Two populations of TrpRSs exist, which bind their amino acid ligand in a distinct way. Two structures from both populations are shown as stick model. Hydrogen bonds (solid blue lines), salt bridges (dashed yellow lines), and hydrophobic interactions (dashed gray lines) are established. A key difference in ligand binding can be observed for a residue that binds the amino group of the indole ring. In human TrpRSs (PDB:1r6u chain A) a hydrogen bond with tyrosine is formed, while *Geobacillus stearothermophilus* (PDB:1i6l chain A) employs aspartic acid.

**Fig. S28.**
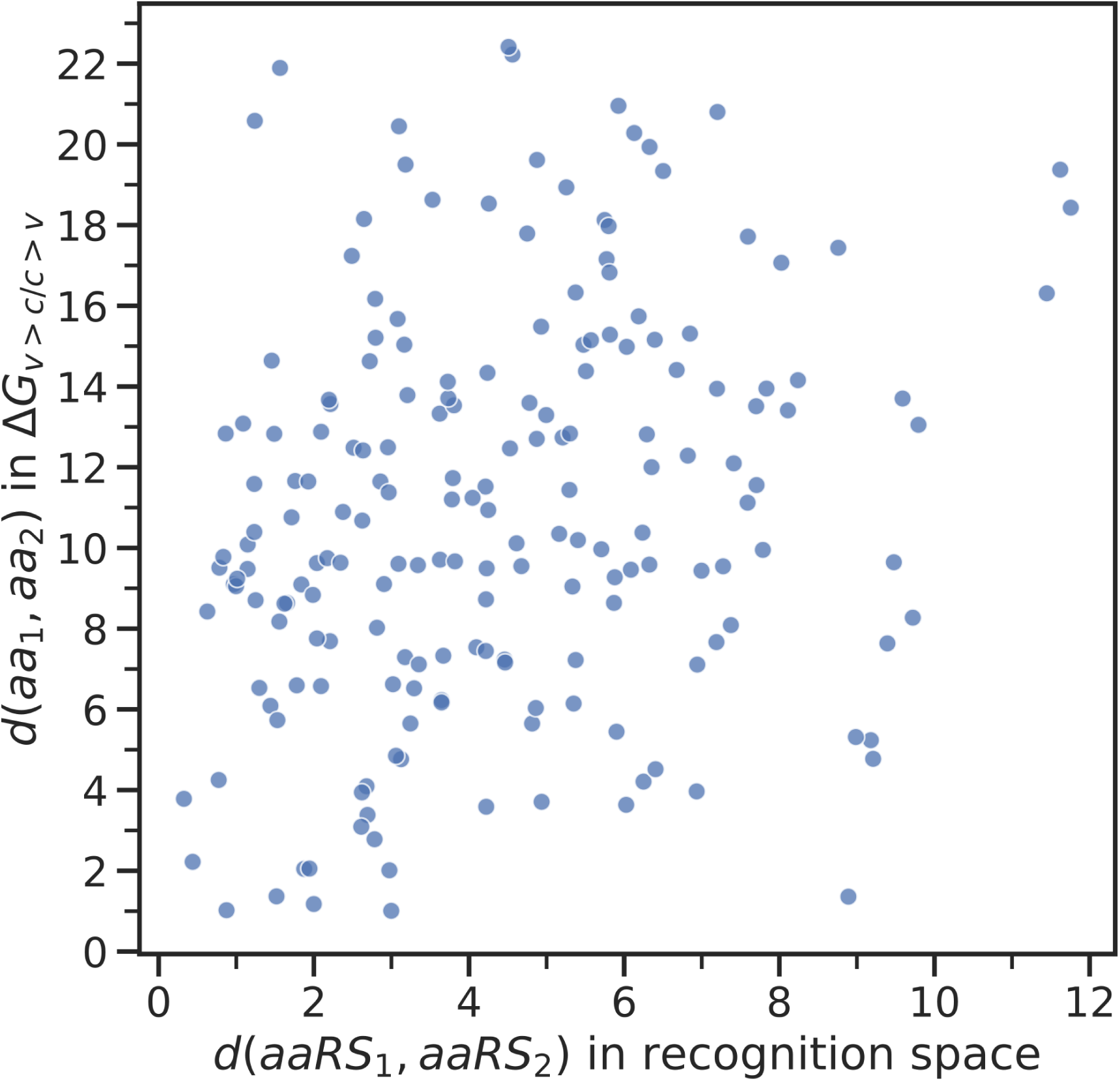
Phase transfer free energies of amino acid side chains (3, 4) from water (Δ*Gw>c*) and vapor (Δ*Gw>c*) to cyclohexane compared to the recognition space analysis from this study. Each data point represents the euclidean distance between every combination of two amino acids in the phase transfer diagram given in Carter and Wills (5) against the euclidean distance in the recognition space proposed in this study. Spearman’s rank correlation is *ρ* = 0.2564 with *p*<0.01.

## References

1. Bernhardt, H. S. The RNA world hypothesis: the worst theory of the early evolution of life (except for all the others)(a). Biol. Direct 7, 23 (2012).

2. Di Giulio, M. The origin of the genetic code: theories and their relationships, a review. BioSystems 80, 175–184 (2005).

3. Carter, C. W. & Wolfenden, R. tRNA acceptor stem and anticodon bases form independent codes related to protein folding. Proc. Natl. Acad. Sci. U.S.A. 112, 7489–7494 (2015).

4. Ibba, M. & Söll, D. Aminoacyl-tRNA synthesis. Annu. Rev. Biochem. 69, 617–650 (2000).

5. Dock-Bregeon, A. et al. Transfer RNA-mediated editing in threonyl-tRNA synthetase. The class II solution to the double discrimination problem. Cell 103, 877–884 (2000).

6. Hadd, A. & Perona, J. J. Coevolution of specificity determinants in eukaryotic glutamyl- and glutaminyl-tRNA synthetases. J. Mol. Biol. 426, 3619–3633 (2014).

7. Nair, N. et al. The Bacillus subtilis and Bacillus halodurans Aspartyl-tRNA Synthetases Retain Recognition of tRNA(Asn). J. Mol. Biol. 428, 618–630 (2016).

8. Song, Y. et al. Double mimicry evades tRNA synthetase editing by toxic vegetable-sourced non-proteinogenic amino acid. Nat Commun 8, 2281 (2017).

9. Carter, C. W. & Wills, P. R. Interdependence, Reflexivity, Fidelity, Impedance Matching, and the Evolution of Genetic Coding. Mol. Biol. Evol. 35, 269–286 (2018).

10. Pak, D., Kim, Y. & Burton, Z. F. Aminoacyl-tRNA synthetase evolution and sectoring of the genetic code. Transcription 1–15 (2018).

11. Swanson, R. et al. Accuracy of in vivo aminoacylation reguires proper balance of tRNA and aminoacyl-tRNA synthetase. Science 242, 1548–1551 (1988).

12. Pham, Y. et al. A Minimal TrpRS Catalytic Domain Supports Sense/Antisense Ancestry of Class I and II Aminoacyl-tRNA Synthetases. Mol. Cell 25, 851–862 (2007).

13. Yu, Y. et al. Crystal structure of human tryptophanyl-tRNA synthetase catalytic fragment: Insights into substrate recognition, tRNA binding, and angiogenesis activity. J. Biol. Chem. 279, 8378–8388 (2004).

14. Doolittle, R. F., Handy, J. & Bada, J. L. Evolutionary anomalies among the aminoacyl-tRNA synthetases. Curr. Opin. Genet. Dev. 8, 630–636 (1998).

15. O’Donoghue, P. & Luthey-Schulten, Z. On the Evolution of Structure in Aminoacyl-tRNA Synthetases. Microbiol. Mol. Biol. Rev. 67, 550–573 (2003).

16. Davis, B. K. Molecular evolution before the origin of species. Prog. Biophys. Mol. Biol. 79, 77–133 (2002).

17. Diaz-Lazcoz, Y. et al. Evolution of genes, evolution of species: the case of aminoacyl-tRNA synthetases. Mol. Biol. Evol. 15, 1548–1561 (1998).

18. Woese, C. R., Olsen, G. J., Ibba, M. & Söll, D. Aminoacyl-tRNA synthetases, the genetic code, and the evolutionary process. Microbiol. Mol. Biol. Rev. 64, 202–236 (2000).

19. Chaliotis, A. et al. The complex evolutionary history of aminoacyl-tRNA synthetases. Nucleic Acids Res. 45, 1059–1068 (2017).

20. Brown, J. R. & Doolittle, W. F. Root of the universal tree of life based on ancient aminoacyl-tRNA synthetase gene duplications. Proc. Natl. Acad. Sci. United States Am. 92, 2441–2445 (1995).

21. Carter, C. W. Coding of Class I and II Aminoacyl-tRNA Synthetases. Adv. Exp. Medicine Biol. 966, 103–148 (2017).

22. Eriani, G., Delarue, M., Poch, O., Gangloff, J. & Moras, D. Partition of tRNA synthetases into two classes based on mutually exclusive sets of sequence motifs. Nature 347, 203–206 (1990).

23. Cusack, S., Berthet-Colominas, C., Hartlein, M., Nassar, N. & Leberman, R. A second class of synthetase structure revealed by X-ray analysis of Escherichia coli seryl-tRNA synthetase at 2.5 A. Nature 347, 249–255 (1990).

24. Cusack, S. Aminoacyl-tRNA synthetases. Curr. Opin. Struct. Biol. 7, 881–889 (1997).

25. Curnow, A. W. et al. Glu-tRNAGln amidotransferase: a novel heterotrimeric enzyme required for correct decoding of glutamine codons during translation. Proc. Natl. Acad. Sci. U.S.A. 94, 11819–11826 (1997).

26. Becker, H. D. & Kern, D. Thermus thermophilus: a link in evolution of the tRNA-dependent amino acid amidation pathways. Proc. Natl. Acad. Sci. U.S.A. 95, 12832–12837 (1998).

27. Hartlein, M. & Cusack, S. Structure, function and evolution of seryl-tRNA synthetases: implications for the evolution of aminoacyl-tRNA synthetases and the genetic code. J. Mol. Evol. 40, 519–530 (1995).

28. Leinfelder, W., Zehelein, E., Mandrand-Berthelot, M. A. & Bock, A. Gene for a novel tRNA species that accepts L-serine and cotranslationally inserts selenocysteine. Nature 331, 723–725 (1988).

29. Sheppard, K. et al. From one amino acid to another: tRNA-dependent amino acid biosynthesis. Nucleic Acids Res. 36, 1813–1825 (2008).

30. Sauerwald, A. et al. RNA-dependent cysteine biosynthesis in archaea. Science 307, 1969–1972 (2005).

31. Rodin, S. N. & Ohno, S. Two types of aminoacyl-tRNA synthetases could be originally encoded by complementary strands of the same nucleic acid. Orig Life Evol Biosph 25, 565–589 (1995).

32. Martinez-Rodriguez, L. et al. Functional Class I and II Amino Acid-activating Enzymes Can Be Coded by Opposite Strands of the Same Gene. J. Biol. Chem. 290, 19710–19725 (2015).

33. Chandrasekaran, S. N., Yardimci, G. G., Erdogan, O., Roach, J. & Carter, C. W. Statistical evaluation of the Rodin-Ohno hypothesis: sense/antisense coding of ancestral class I and II aminoacyl-tRNA synthetases. Mol. Biol. Evol. 30, 1588–1604 (2013).

34. Carter, C. W. Urzymology: experimental access to a key transition in the appearance of enzymes. J. Biol. Chem. 289, 30213–30220 (2014).

35. Carter, C. W. et al. The Rodin-Ohno hypothesis that two enzyme superfamilies descended from one ancestral gene: an unlikely scenario for the origins of translation that will not be dismissed. Biol. Direct 9, 11 (2014).

36. Arnez, J. G. & Moras, D. Structural and functional considerations of the aminoacylation reaction. Trends Biochem. Sci. 22, 211–216 (1997).

37. Praetorius-Ibba, M. et al. Ancient adaptation of the active site of tryptophanyl-tRNA synthetase for tryptophan binding. Biochemistry 39, 13136–13143 (2000).

38. Kaiser, F. et al. Backbone Brackets and Arginine Tweezers delineate Class I and Class II aminoacyl tRNA synthetases. PLoS Comput. Biol. 14, e1006101 (2018).

39. Klebe, G. & Bohm, H. J. Energetic and entropic factors determining binding affinity in protein-ligand complexes. J. Recept. Signal Transduct. Res. 17, 459–473 (1997).

40. Salentin, S., Haupt, V. J., Daminelli, S. & Schroeder, M. Polypharmacology rescored: protein-ligand interaction profiles for remote binding site similarity assessment. Prog. Biophys. Mol. Biol. 116, 174–186 (2014).

41. Salentin, S., Schreiber, S., Haupt, V. J., Adasme, M. F. & Schroeder, M. PLIP: fully automated protein-ligand interaction profiler. Nucleic Acids Res. 43, W443–447 (2015).

42. Berman, H. M. et al. The Protein Data Bank. Nucleic Acids Res. 28, 235–242 (2000).

43. Livingstone, C. D. & Barton, G. J. Protein sequence alignments: a strategy for the hierarchical analysis of residue conservation. Comput. Appl. Biosci. 9, 745–756 (1993).

44. Bonnici, V., Giugno, R., Pulvirenti, A., Shasha, D. & Ferro, A. A subgraph isomorphism algorithm and its application to biochemical data. BMC Bioinforma. 14 Suppl 7, S13 (2013).

45. Perona, J. J. & Gruic-Sovulj, I. Synthetic and editing mechanisms of aminoacyl-tRNA synthetases. Top Curr Chem 344, 1–41 (2014).

46. Fersht, A. R. et al. Hydrogen bonding and biological specificity analysed by protein engineering. Nature 314, 235–238 (1985).

47. Fukai, S. et al. Structural basis for double-sieve discrimination of L-valine from L-isoleucine and L-threonine by the complex of tRNA(Val) and valyl-tRNA synthetase. Cell 103, 793–803 (2000).

48. Zivkovic, I., Moschner, J., Koksch, B. & Gruic-Sovulj, I. Mechanism of discrimination of isoleucyl-tRNA synthetase against nonproteinogenic Î±-aminobutyrate and its fluorinated analogues. FEBS J. (2019).

49. Guo, M. et al. Paradox of mistranslation of serine for alanine caused by AlaRS recognition dilemma. Nature 462, 808–812 (2009).

50. Fersht, A. R. & Dingwall, C. Evidence for the Double-Sieve Editing Mechanism in Protein Synthesis. Steric Exclusion of Isoleucine by Valyl-tRNA Synthetases. Biochemistry 18, 2627–2631 (1979).

51. Kaiser, F., Eisold, A., Bittrich, S. & Labudde, D. Fit3D: a web application for highly accurate screening of spatial residue patterns in protein structure data. Bioinformatics 32, 792–794 (2016).

52. Durrant, J. D., Votapka, L., Sørensen, J. & Amaro, R. E. POVME 2.0: An Enhanced Tool for Determining Pocket Shape and Volume Characteristics. J Chem Theory Comput. 10, 5047–5056 (2014).

53. Crooks, G. E., Hon, G., Chandonia, J. M. & Brenner, S. E. WebLogo: a sequence logo generator. Genome Res. 14, 1188–1190 (2004).

54. Schrödinger, LLC. The PyMOL molecular graphics system, version 1.8 (2015).

55. McInnes, L. & Healy, J. UMAP: Uniform Manifold Approximation and Projection for Dimension Reduction. ArXiv e-prints (2018).

56. Rousseeuw, P. J. Silhouettes: A graphical aid to the interpretation and validation of cluster analysis. J. Comput. Appl. Math. 20, 53–65 (1987).

57. Wolfenden, R., Lewis, C. A., Yuan, Y. & Carter, C. W. Temperature dependence of amino acid hydrophobicities. Proc. Natl. Acad. Sci. U.S.A. 112, 7484–7488 (2015).

58. W. Carter, C. & Wills, P. Did gene expression co-evolve with gene replication? In Evolutionary Biology: Origin and Evolution of Biodiversity, 293–313, DOI: 10.1007/978-3-319-95954-2_16 (Springer International Publishing, 2018).

59. Dutta, S., Choudhury, K., Banik, S. D. & Nandi, N. Active site nanospace of aminoacyl tRNA synthetase: difference between the class I and class II synthetases. J Nanosci Nanotechnol 14, 2280–2298 (2014).

60. Davis, B. K. Evolution of the genetic code. Prog. Biophys. Mol. Biol. 72, 157–243 (1999).

61. Wong, J. T. F. A co-evolution theory of the genetic code. Proc. Natl. Acad. Sci. 72, 1909–1912 (1975).

62. Klipcan, L. & Safro, M. Amino acid biogenesis, evolution of the genetic code and aminoacyl-tRNA synthetases. J. Theor. Biol. 228, 389–396 (2004).

63. Weber, A. L. & Miller, S. L. Reasons for the occurrence of the twenty coded protein amino acids. J. Mol. Evol. 17, 273–284 (1981).

64. Rogers, S. O. Evolution of the genetic code based on conservative changes of codons, amino acids, and aminoacyl tRNA synthetases. J. Theor. Biol. 466, 1–10 (2019).

65. Newton, M. S., Morrone, D. J., Lee, K. H. & Seelig, B. Genetic Code Evolution Investigated through the Synthesis and Characterisation of Proteins from Reduced-Alphabet Libraries. Chembiochem 20, 846–856 (2019).

66. Cammer, S. & Carter, C. W. Six Rossmannoid folds, including the Class I aminoacyl-tRNA synthetases, share a partial core with the anti-codon-binding domain of a Class II aminoacyl-tRNA synthetase. Bioinformatics 26, 709–714 (2010).

67. Bittrich, S. et al. Application of an interpretable classification model on Early Folding Residues during protein folding. BioData Min 12, 1 (2019).

68. Lu, M. F., Xie, Y., Zhang, Y. J. & Xing, X. Y. Effects of Cofactors on Conformation Transition of Random Peptides Consisting of a Reduced Amino Acid Alphabet. Protein Pept. Lett. 22, 579–585 (2015).

69. Kang, S. K. et al. ATP selection in a random peptide library consisting of prebiotic amino acids. Biochem. Biophys. Res. Commun. 466, 400–405 (2015).

70. Wong, J. T. F. Coevolutlon theory of genetic code at age thirty. BioEssays 27, 416–425 (2005).

71. Griffiths, G. Cell evolution and the problem of membrane topology. Nat. Rev. Mol. Cell Biol. 8, 1018–1024 (2007).

72. Brooks, D. J., Fresco, J. R., Lesk, A. M. & Singh, M. Evolution of amino acid frequencies in proteins over deep time: inferred order of introduction of amino acids into the genetic code. Mol. Biol. Evol. 19, 1645–1655 (2002).

73. Trifonov, E. N. The triplet code from first principles. J. Biomol. Struct. Dyn. 22, 1–11 (2004).

74. Martinis, S. A. & Boniecki, M. T. The balance between pre- and post-transfer editing in tRNA synthetases. FEBS Lett. 584, 455–459 (2010).

75. Schulze, J. O. et al. Crystal structure of a non-discriminating glutamyl-tRNA synthetase. J. Mol. Biol. 361, 888–897 (2006).

76. Perona, J. J., Rould, M. A. & Steitz, T. A. Structural Basis for Transfer RNA Aminoacylation by Escherichia coli Glutaminyl-tRNA Synthetase. Biochemistry 32, 8758–8771 (1993).

77. Bullock, T. L., Uter, N., Nissan, T. A. & Perona, J. J. Amino acid discrimination by a class I aminoacyl-tRNA synthetase specified by negative determinants. J. Mol. Biol. 328, 395–408 (2003).

78. Sever, S., Rogers, K., Rogers, M. J., Carter, C. & Söll, D. Escherichia coli tryptophanyl-tRNA synthetase mutants selected for tryptophan auxotrophy implicate the dimer interface in optimizing amino acid binding. Biochemistry 35, 32–40 (1996).

79. Iwasaki, W. et al. Structural Basis of the Water-assisted Asparagine Recognition by Asparaginyl-tRNA Synthetase. J. Mol. Biol. 360, 329–342 (2006).

80. Jakubowski, H. Misacylation of tRNA(Lys) with noncognate amino acids by Lysyl-tRNA synthetase. Biochemistry 38, 8088–8093 (1999).

81. Nagel, G. M. & Doolittle, R. F. Evolution and relatedness in two aminoacyl-tRNA synthetase families. Proc. Natl. Acad. Sci. United States Am. 88, 8121–8125 (1991).

82. Qin, X. et al. Cocrystal structures of glycyl-tRNA synthetase in complex with tRNA suggest multiple conformational states in glycylation. J. Biol. Chem. 289, 20359–20369 (2014).

83. Naganuma, M., Sekine, S., Fukunaga, R. & Yokoyama, S. Unique protein architecture of alanyl-tRNA synthetase for aminoacylation, editing, and dimerization. Proc. Natl. Acad. Sci. U.S.A. 106, 8489–8494 (2009).

84. Nakayama, T. et al. Deficient activity of alanyl-tRNA synthetase underlies an autosomal recessive syndrome of progressive microcephaly, hypomyelination, and epileptic encephalopathy. Hum. Mutat. 38, 1348–1354 (2017).

85. Splan, K. E., Ignatov, M. E. & Musier-Forsyth, K. Transfer RNA modulates the editing mechanism used by class II prolyl-tRNA synthetase. J. Biol. Chem. 283, 7128–7134 (2008).

86. Rayevsky, A., Sharifi, M. & Tukalo, M. A molecular dynamics simulation study of amino acid selectivity of LeuRS editing domain from Thermus thermophilus. J. Mol. Graph. Model. 84, 74–81 (2018).

87. Doublie, S., Bricogne, G., Gilmore, C. & Carter, C. W. Tryptophanyl-tRNA synthetase crystal structure reveals an unexpected homology to tyrosyl-tRNA synthetase. Structure 3, 17–31 (1995).

88. Wolf, Y. I., Aravind, L., Grishin, N. V. & Koonin, E. V. Evolution of Aminoacyl-tRNA Synthetases -Analysis of Unique Domain Architectures and Phylogenetic Trees Reveals a Complex History of Horizontal Gene Transfer Events. Genome Res. 9, 689–710 (1999).

89. Fournier, G. P. & Alm, E. J. Ancestral Reconstruction of a Pre-LUCA Aminoacyl-tRNA Synthetase Ancestor Supports the Late Addition of Trp to the Genetic Code. J. Mol. Evol. 80, 171–185 (2015).

90. Weinreb, V. et al. Enhanced amino acid selection in fully evolved tryptophanyl-tRNA synthetase, relative to its urzyme, requires domain motion sensed by the D1 switch, a remote dynamic packing motif. J. Biol. Chem. 289, 4367–4376 (2014).

91. Li, L. & Carter, C. W. Full implementation of the genetic code by tryptophanyl-tRNA synthetase requires intermodular coupling. J. Biol. Chem. 288, 34736–34745 (2013).

92. Ribas de Pouplana, L., Frugier, M., Quinn, C. L. & Schimmel, P. Evidence that two present-day components needed for the genetic code appeared after nucleated cells separated from eubacteria. Proc. Natl. Acad. Sci. U.S.A. 93, 166–170 (1996).

93. Yang, X. L. et al. Crystal structures that suggest late development of genetic code components for differentiating aromatic side chains. Proc. Natl. Acad. Sci. U.S.A. 100, 15376–15380 (2003).

94. Needleman, S. B. & Wunsch, C. D. A general method applicable to the search for similarities in the amino acid sequence of two proteins. J. Mol. Biol. 48, 443–453 (1970).

95. Prlic, A. et al. BioJava: an open-source framework for bioinformatics in 2012. Bioinformatics 28, 2693–2695 (2012).

96. Notredame, C., Higgins, D. G. & Heringa, J. T-Coffee: A novel method for fast and accurate multiple sequence alignment. J. Mol. Biol. 302, 205–217 (2000).

97. Leberecht, C., Kaiser, F., Bittrich, S. & Krautwurst, S. cleberecht/singa: singa-all release v0.4.0, DOI: 10.5281/zenodo. 1320146 (2018).

98. Kim, S. et al. PubChem Substance and Compound databases. Nucleic Acids Res. 44, D1202–1213 (2016)

## References

1. G. F Kaiser, et al., Backbone Brackets and Arginine Tweezers delineate Class I and Class II aminoacyl tRNA synthetases. PLoS Comput. Biol. 14, e1006101 (2018).

2. H. GE Crooks, G Hon, JM Chandonia, SE Brenner, WebLogo: a sequence logo generator. Genome Res. 14, 1188–1190 (2004).

3. I. CW Carter, R Wolfenden, tRNA acceptor stem and anticodon bases form independent codes related to protein folding. Proc. Natl. Acad. Sci. U.S.A. 112, 7489–7494 (2015).

4. J. R Wolfenden, CA Lewis, Y Yuan, CW Carter, Temperature dependence of amino acid hydrophobicities. Proc. Natl. Acad. Sci. U.S.A. 112, 7484–7488 (2015).

5. K. C W. Carter, P Wills, Did Gene Expression Co-evolve with Gene Replication? (Springer International Publishing), pp. 293–313 (2018).

